# Natural leaf shape variation reveals diverse transcriptional targets of *GmJAG1* during soybean leaf development

**DOI:** 10.64898/2026.04.08.717315

**Authors:** Bishal G. Tamang, Casey Kramer, Elizabeth A. Ainsworth

## Abstract

The *JAGGED* transcription factor family regulates lateral organ development across angiosperms. In soybean (*Glycine max* Merr.), a D9H mutation in the EAR repression motif of *GmJAG1* causes a narrow leaflet phenotype and explains over 70% of phenotypic variance in leaf shape. Because this mutation does not affect the zinc finger DNA-binding domain, both alleles bind identical targets but differ in repressor recruitment. Previous studies mapped *GmJAG1* binding sites, but the functional targets controlling leaf morphology are uncharacterized. Here, we used comparative transcriptomics across four soybean genotypes with contrasting leaf shape, spanning a developmental time series from shoot apex to mature leaf, and identified 1,567 candidate target genes. *GmJAG1* expression was confined to the shoot apex, yet 99.1% of candidate targets maintained differential expression throughout development. We found that neither Kip-Related Protein (*KRP*) cell cycle inhibitors nor Cyclin-Dependent Kinases (CDKs) showed differential expression despite binding evidence in Arabidopsis. However, D-type cyclins were upregulated in narrow-leaf genotypes suggesting cyclin-mediated rather than *KRP*-mediated cell cycle regulation in soybean. Pathway analysis revealed enrichment of auxin (1.8-fold, P = 0.02) and salicylic acid (4-fold, P = 0.016) genes among JAG1_D9H_ targets. Filtering by differential expression, binding data, phenotype correlation, and co-expression network membership identified 79 high-confidence targets, including orthologs of *NPH3* (phototropin-mediated leaf flattening), *MIK2* (cell wall integrity sensing), *RD22* (ABA-responsive stress signaling), and *SCL23* (GRAS transcription factor in bundle sheath development). These candidates provide targets for functional validation and breeding in legumes.

## Introduction

Leaf morphology influences light interception, gas exchange, photosynthesis, and ultimately crop yield (Zhu et al., 2010). In soybean (*Glycine max* Merr.), canopy architecture has been linked to photosynthetic efficiency and yield potential, with historical yield gains driven by improvements in light interception efficiency, energy conversion, and harvest index (Koester et al., 2014). Modern soybean cultivars often have higher than optimal leaf area index (LAI) which can limit light penetration into lower canopy layers and reduce canopy photosynthesis (Slattery and Ort, 2021). There is therefore growing interest in the genetic basis of leaf shape variation to optimize canopy architecture. Recent field studies showed that introgression of the narrow leaflet allele (*ln*) reduces LAI by ∼13% without yield penalty (Tamang et al., 2026a) which suggests that modifying leaf shape could be one approach to canopy optimization.

The genetics of soybean leaf shape have been studied for over a century, with the first reports of the narrow leaflet trait dating to Takahashi and Fukuyama (1919). The *Ln* locus controls the transition between broad (*Ln*) and narrow (*ln*) leaflet phenotypes with the narrow allele inherited in an incompletely dominant manner (Bernard and Weiss, 1973). In near-isogenic backgrounds, this locus explains over 70% of the variation in leaf shape (Tamang et al., 2026c), making it one of the largest-effect loci for leaf shape in any crop. Fine genetic mapping localized the causal variant to a 12.6-kb region on chromosome 20 containing *GmJAG1* (*Glyma.20G116200*), a C2H2 zinc finger transcription factor homologous to Arabidopsis *JAGGED* (Jeong et al., 2012). The soybean genome retains ∼75% of its genes in duplicate from two whole-genome duplication events (Schmutz et al., 2010), and the synteny between chromosomes 20 and 10 gave rise to the *GmJAG2* paralog (*Glyma.10G273800*) on chromosome 10. The narrow leaflet phenotype is caused by a D9H amino acid substitution in the conserved EAR (ethylene-responsive element binding factor-associated amphiphilic repression) motif of the *GmJAG1* protein. This mutation affects the repression domain rather than the zinc finger DNA-binding domain, so both protein variants presumably bind the same targets but differ in co-repressor recruitment. More recently, *GmJAG1* DNA-binding preferences have been characterized using ChIPmentation on soybean protoplasts (Huang et al., 2021) and DAP-seq with recombinant proteins (Wang et al., 2024). These studies identified thousands of potential binding sites across the genome.

*JAGGED* (*JAG*) transcription factors are conserved regulators of lateral organ development. In Arabidopsis, *JAG* promotes tissue growth by suppressing premature differentiation in developing organs (Dinneny et al., 2004; Ohno et al., 2004; Bar and Ori, 2014), with the paralog *NUBBIN* (*NUB*) acting redundantly in reproductive organs (Dinneny et al., 2006). *JAG* family proteins contain an EAR motif (LDLNNLP in *GmJAG1*) that recruits TOPLESS (*TPL*) and TOPLESS-RELATED (*TPR*) co-repressors and histone deacetylases to establish repressive chromatin at target loci (Szemenyei et al., 2008; Kagale and Rozwadowski, 2011; Causier et al., 2012). The D9H mutation in soybean *GmJAG1* disrupts a conserved aspartate of the EAR motif (Jeong et al., 2012), potentially compromising *TPL*/*TPR* recruitment and repression function.

The *JAG* molecular mechanism was first characterized in Arabidopsis, where it controls the transition from isotropic to anisotropic growth and overrides the G1/S cell-size checkpoint (Schiessl et al., 2012). *JAG* directly represses KIP-RELATED PROTEIN (*KRP*) cell cycle inhibitors, particularly *KRP2* and *KRP4*, which bind and inhibit CDK-cyclin complexes (De Veylder et al., 2001; Inzé and De Veylder, 2006; Schiessl et al., 2014). Genetic evidence confirmed this as *krp2* and *krp4* mutations partially suppressed *jag* growth defects (Schiessl et al., 2014). This established a "double-negative" regulatory logic where *JAG* represses inhibitors, thereby activating cell cycle progression. Schiessl et al. (2014) also identified D-type cyclin (*CYCD3;3*) among *JAG*-repressed targets, though this was not explored further.

D-type cyclins (CYCDs) bind CDKs to form active kinase complexes that drive the G1/S transition (Dewitte et al., 2003; Menges et al., 2006). In Arabidopsis, *CYCD3* overexpression causes hyperproliferation of cells and altered leaf architecture (Dewitte et al., 2003, 2007), and SQUAMOSA PROMOTER BINDING PROTEIN-LIKE 9 (*SPL9*) activates *CYCD3* to encode developmental timing controlling leaf morphogenesis (Li et al., 2024), establishing cyclins as effectors linking developmental programs to leaf shape. In soybean, *GmPLATZ* directly activates *CYCD1;1* and *CYCD6;1* to promote seed size through cell proliferation rather than cell expansion (Hu et al., 2023), confirming the functional importance of D-type cyclins in this species.

While *JAG* function is well characterized in Arabidopsis, several questions remain about its role in legumes and crops like soybean. Although the *ln* locus in soybean has been mapped to *GmJAG1* and the D9H mutation has been identified, the genome-wide targets of *GmJAG1* have not been systematically characterized. Recent binding studies identified thousands of potential *GmJAG1* binding sites (Huang et al., 2021; Wang et al., 2024). However, transcription factor binding does not necessarily equate to transcriptional regulation, and the functional significance of these binding events remains to be tested. Given the evolutionary distance between Arabidopsis and soybean (∼90 million years; Grant et al., 2000), it is also unclear whether the *JAG-KRP* regulatory connection is conserved. Additionally, the narrow leaflet phenotype is associated with anatomical differences including increased leaf thickness (Tamang et al., 2023) which suggests that *JAG1* may regulate developmental processes beyond cell proliferation. Whether *GmJAG1* candidate targets include hormone signaling pathway genes, as might be expected from EAR domain function in other contexts, has not been examined. Therefore, understanding the *GmJAG1* regulatory network would help clarify how leaf shape is determined in legumes and could inform breeding strategies for canopy optimization.

To address these questions, we used natural leaf shape variation as an *in vivo* system to identify candidate *GmJAG1* targets. By comparing transcriptomes of soybean genotypes with functional versus non-functional *GmJAG1* across a developmental time series, we can identify genes whose expression changes when *JAG1* function is disrupted. This complements binding studies that show where *JAG1* binds but cannot establish whether that binding leads to changes in gene expression. We had four objectives: (1) identify high-confidence genome-wide transcriptional targets of *GmJAG1* using a tiered classification based on reproducibility across genetic backgrounds; (2) integrate differential expression with published ChIP-seq and DAP-seq binding data, phenotypic correlation and co-expression networks to identify the highest-confidence direct targets; (3) characterize the temporal dynamics of *JAG1*-regulated genes across development; and (4) test whether the *JAG1-KRP* regulatory relationship from Arabidopsis is conserved in soybean. We identified 1,567 candidate target genes with 79 supported by all evidence types, including orthologs of genes controlling cell proliferation, cell wall integrity and hormone homeostasis in Arabidopsis. Additionally, *KRP* cell cycle inhibitors showed no transcriptional response despite binding evidence in Arabidopsis, while D-type cyclins were regulated, suggesting divergent cell cycle control mechanisms in soybean.

## Results

### Broad and Narrow Genotypes Show Progressive Leaf Divergence

We performed bulk RNA-seq analysis on leaves of four soybean genotypes with contrasting leaf morphology at five developmental timepoints (TP1-TP5) spanning shoot apex (TP1) through mature leaf expansion (TP5) (Figure 1A). In chamber-grown plants, the two narrow-leaved genotypes (PI 547745 and PI 612713B) had leaf length-to-width (L:W) ratios of 2.16 and 2.36, while two broad-leaved genotypes (LD11-2170 and PI 532462A) showed L:W ratios of 1.59 and 1.55, respectively (Figure 1A).

**Figure 1.**
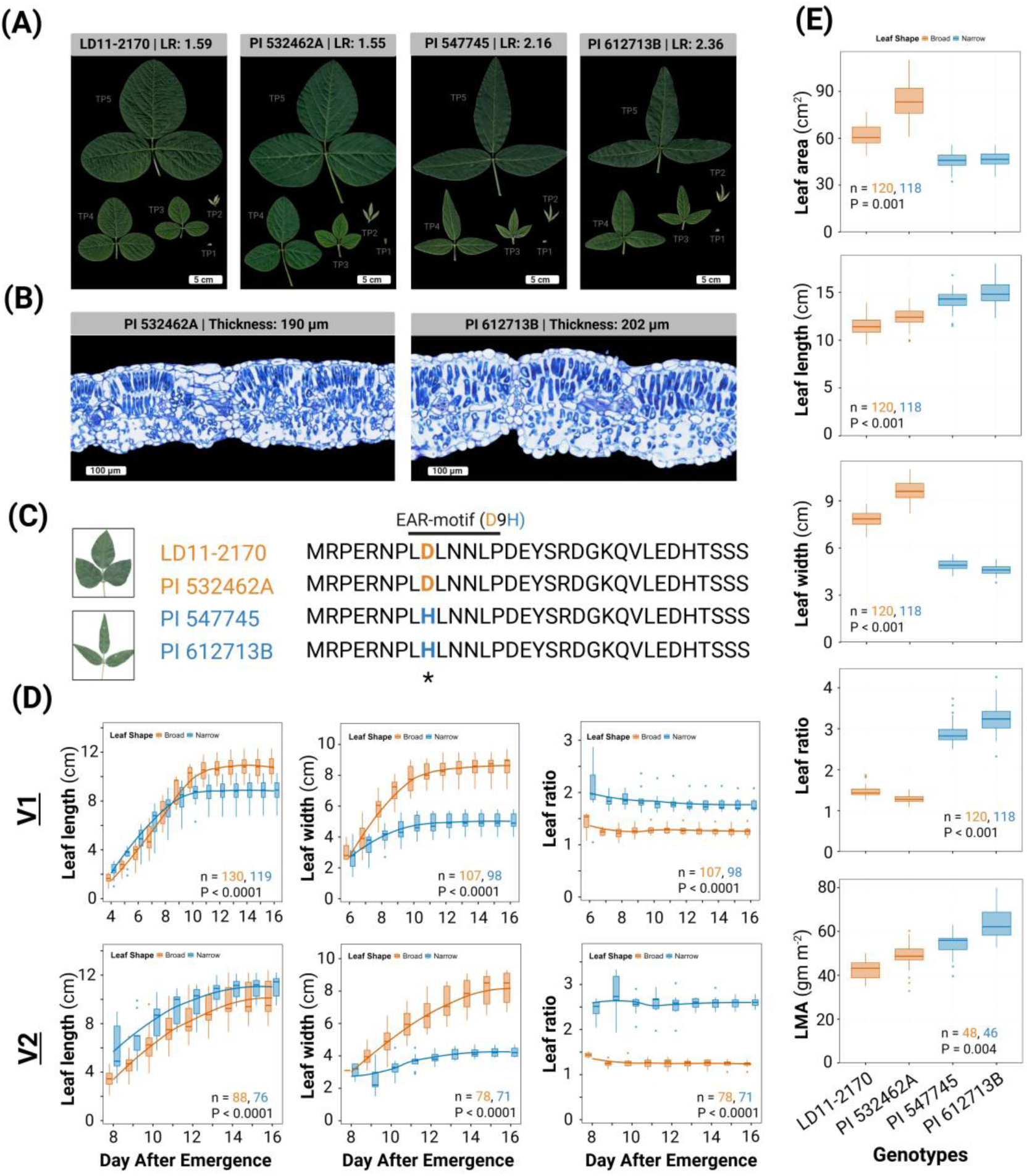
Phenotypic characterization of broad and narrow leaf soybean genotypes. **(A)** Representative trifoliate leaf images across five developmental stages (TP1-TP5) corresponding to RNA-seq tissue collection time points. **(B)** Leaf cross-sections stained with toluidine blue for PI 532462A (left) and PI 612713B (right). **(C)** Amino acid alignment of the N-terminal 30 residues of GmJAG1 across all four genotypes. The EAR repression motif (LDLNNLP, positions 8-14) is annotated. An asterisk (*) marks the mutation site. **(D)** Temporal dynamics of leaf morphological traits in broad and narrow leaf genotypes. Rows correspond to leaf nodes: V1 middle trifoliate (top) and V2 middle trifoliate (bottom). Columns show leaf length (left), leaf width (middle), and length-to-width ratio (right). Box-and-whisker plots show the distribution at each time point with LOESS-smoothed trend lines. Sample sizes (n) and two-way ANOVA P-values for the leaf type main effect are shown within each panel. **(E)** Field-grown leaf morphological traits and leaf mass per area (LMA) for the same four genotypes. P-values are from RCBD ANOVA for the leaf type main effect.

Leaf cross-sections from three of the four genotypes showed mean thickness of 190 ± 13 *µ*m for PI 532462A (broad; n = 3), 203 ± 10 *µ*m for PI 547745 (narrow; n = 3) and 202 ± 11 *µ*m for PI 612713B (narrow; n = 3) (Figure 1B). This thickness did not differ significantly among these three genotypes (one-way ANOVA, P = 0.4). The two narrow-leaved genotypes (PI 547745 and PI 612713B) carried the D9H substitution in the *GmJAG1* EAR motif, while the two broad-leaved genotypes (LD11-2170 and PI 532462A) retained the functional aspartate (Figure 1C).

The leaf shape divergence between broad- and narrow-leaved genotypes was evident from the earliest measurement timepoint. Narrow leaves showed approximately 1.5-fold higher L:W ratios at the V1 node (1.77 vs. 1.26) and 2-fold higher ratios at the V2 node (2.53 vs. 1.24) compared to broad leaves (Figure 1D). Two-way ANOVA (Leaf type x Day) revealed that absolute leaf dimensions diverged progressively, with significant Leaf x Day interactions for width at V1 (P = 4.2 x 10^-^¹⁵) and V2 (P = 2.8 x 10^-^⁸) nodes. In contrast, the length-to-width ratio showed no significant Leaf x Day interaction at any node (all P > 0.2).

Field measurements confirmed the leaf shape differences observed in the controlled environment. Broad-leaved genotypes had significantly greater leaf area (73.6 ±14.7 vs. 46.3 ± 4.7 cm²; P = 0.001) and maximum width (8.8 ± 1.1 vs. 4.8 ± 0.3 cm; P < 0.001), while narrow-leaved genotypes had greater length (14.6 ± 1.2 vs. 11.9 ± 1.0 cm; P < 0.001) and higher length-to-width ratios (3.08 ± 0.35 vs. 1.37 ± 0.14; P < 0.001) (Figure 1E). Narrow-leaved genotypes also had significantly higher LMA (58.9 ± 8.4 vs. 45.5 ± 6.3 g m^-^²; P = 0.004).

### Transcriptome Profiling Reveals Clear Separation between Leaf Developmental Stages

Three biological replicates per genotype-timepoint combination were collected yielding a total of 60 samples for transcriptome analysis. RNA sequencing generated 1.77 billion reads across all 60 samples, with library sizes ranging from 16.9 to 46.3 million reads per sample (Figure S1A). After filtering genes with low expression (CPM > 1 in at least 3 samples), 35,941 genes (74.3% of 48,359 initial genes) were retained for downstream analysis. The filtered-out genes contributed minimally to read counts (99.9% of reads were retained).

ComBat-seq batch correction was applied to address batch effects between batch 1 and batch 2 sequencing runs. PVCA showed that batch-associated variance decreased from 34.6% to 8.0% while biological variance (timepoint + leaf type) increased from 61.6% to 85.1% (Figure 2A; Figure S1B-D). After correction, sample correlations showed high reproducibility, with mean within-replicate correlations ranging from 0.87 to 0.98 (Figure 2B).

**Figure 2.**
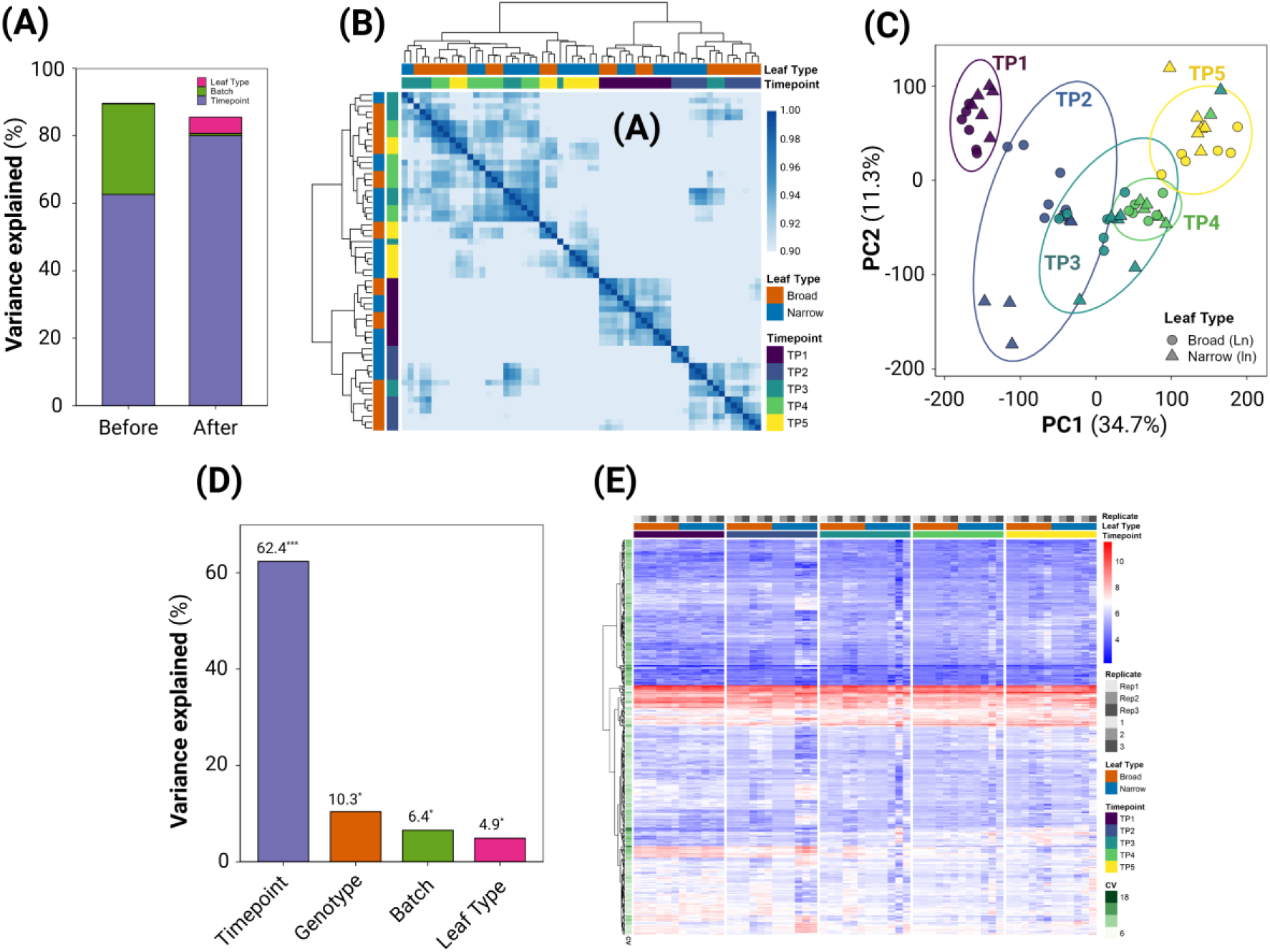
RNA-seq data quality assessment and validation. **(A)** Principal Variance Component Analysis (PVCA) showing variance decomposition before and after ComBat-seq batch correction. Stacked bar plot displays the proportion of variance explained by Timepoint, Batch, and Leaf Type. **(B)** Sample correlation heatmap showing Pearson correlation coefficients across all 60 samples. Annotation bars indicate timepoint (purple-yellow gradient), leaf type (orange: Broad; blue: Narrow), and genotype. **(C)** Principal component analysis (PCA) of batch-corrected, TMM-normalized gene expression data. 95% confidence ellipses are drawn around each timepoint group. **(D)** PERMANOVA analysis quantifying variance explained (R²) by experimental factors. Significance: *** p < 0.001. **(E)** Expression stability of housekeeping genes across all samples. Heatmap displays log_2_CPM values for reference genes from Machado et al. (2020). Genes are clustered by row; samples are ordered by timepoint (TP1-TP5) with gaps between timepoints. Coefficient of variation (CV) is shown as row annotation.

Similarly, PCA revealed that PC1 explained 34.7% of variance and PC2 explained 11.3% with clear separation of samples by developmental timepoint along PC1 (Figure 2C). PERMANOVA analysis confirmed that timepoint explained the largest proportion of variance (R^2^ = 62.4%, p < 0.001) followed by genotype (R^2^ = 10.4%, p = 0.029), batch (R^2^ = 6.5%, p = 0.013) and leaf type (R^2^ = 4.9%, p = 0.029) (Figure 2D). Established soybean housekeeping genes showed stable expression (median CV = 7.7% vs. 16.4% genome-wide) confirming dataset quality (Figure 2E; Table S1).

### Differential Expression Analysis Identified 1,567 GmJAG1 Candidate Genes

*GmJAG1* expression showed a distinct temporal pattern where expression was observed only at TP1 (mean log_2_CPM = 0.85; Figure 3A). It declined to near-undetectable levels from TP2 onward (mean log_2_CPM < -5). The *JAG2* paralog (*Glyma.10G273800*) showed a similar pattern, with expression confined to TP1 as well (broad: 1.82 log_2_CPM and narrow: 1.92 log_2_CPM) and declining to near-zero by TP2. Expression levels of both *JAG*s did not differ significantly between leaf types at TP1 (P > 0.05). qRT-PCR independently confirmed expression of both *GmJAG1* and *GmJAG2* at TP1 in all four genotypes, with no significant difference between leaf types for either gene (*JAG1* P = 0.53; *JAG2* P = 0.91; Figure 3A).

**Figure 3.**
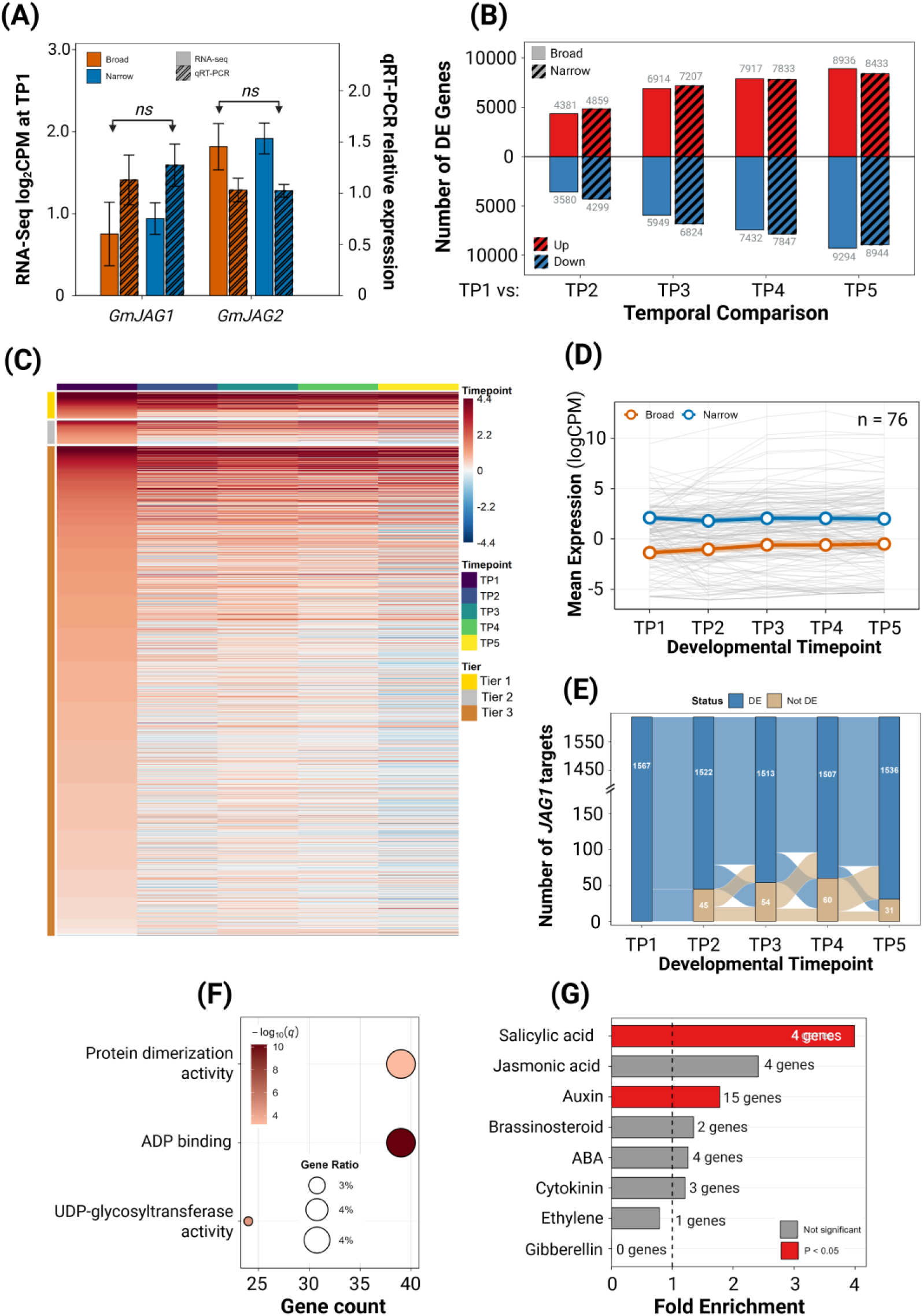
GmJAG1 target identification and temporal dynamics. **(A**) Expression of GmJAG1 and GmJAG2 at the shoot apex (TP1). Solid bars represent RNA-seq expression (left y-axis, mean log₂CPM); striped bars represent qRT-PCR relative expression (right y-axis, n = 6 per leaf type). Error bars indicate standard error. **(B)** Temporal differential expression analysis comparing each developmental stage to TP1. **(C)** Temporal expression heatmap of 1,567 differentially expressed genes across developmental timepoints (TP1-TP5). Rows represent individual genes grouped by confidence tier (Tier 1, Tier 2, Tier 3); columns represent timepoints. Genes are ordered by TP1 fold change magnitude within each tier. **(D)** Mean expression trajectories of Tier 1 DEGs across developmental timepoints. Individual gene trajectories shown in grey; mean expression ± SE shown for Broad and Narrow genotypes. (**E)** Alluvial plot showing DE status of 1,567 DEGs across developmental timepoints (TP1-TP5). Bar height indicates number of genes classified as differentially expressed (DE, blue) or not differentially expressed (Not DE, tan) at each timepoint. Flows between bars illustrate gene transitions between states. Y-axis contains a scale break between 150 and 1,400. **(F)** Gene Ontology enrichment analysis of 1,567 DEGs. Dot plot shows significantly enriched GO terms (q < 0.05) from Molecular Function (MF) categories. Point size indicates gene ratio; color intensity represents statistical significance (-log_10_q-value). **(G)** Hormone pathway enrichment among JAG1 candidate targets. Bar chart shows fold enrichment of hormone-related genes in the DEG set compared to genome background. Red bars indicate significantly enriched pathways (P < 0.05); dashed line represents expected fold enrichment (1.0).

Temporal differential expression analysis revealed a progressive transcriptional change from the shoot apex stage, with TP4 and TP5 showing the most differentially expressed genes (DEGs) relative to TP1 in both broad and narrow leaves (Figure 3B). Differential expression analysis comparing broad and narrow-leaved genotypes included four pairwise comparisons (each narrow versus broad genotype) and one pooled comparison (all narrow versus all broad genotypes). Using thresholds of FDR < 0.05 and |log_2_FC| > 1, the pooled comparison identified 1,098 DEGs (721 upregulated representing 65.7% and 377 downregulated in narrow-leaved genotypes) (Table S3).

The tiered classification system (defined in the Materials and Methods section) based on reproducibility across pairwise comparisons identified 1,567 putative*JAG1* target genes (Figure 3C; Table S2). Genes differentially expressed in all four pairwise comparisons were classified as Tier 1 (76 genes), those in three of four pairwise comparisons as Tier 2 (69 genes) and those in two of four pairwise comparisons or the pooled comparison as Tier 3 (1,422 genes). Tier 1 targets showed the highest mean fold changes (mean |log_2_FC| = 3.8) (Table S2). These log_2_ fold changes ranged between 1.2 and 12.2, with five genes showing log_2_FC > 10. Among Tier 1 targets, 74 of 76 genes (97.4%) showed coordinate expression trajectories with clear separation between broad and narrow genotypes maintained across all timepoints (Figure 3D).

Despite *GmJAG1* expression being confined to TP1, 1,553 of 1,567 candidate targets (99.1%) maintained differential expression from shoot apex through mature leaf development, with only 14 showing early transient patterns (Figure 3E; Figure S2A). Tracking DE status across all five timepoints showed that >95% of targets remained differentially expressed at each stage.

### GO Enrichment Identifies ADP Binding, Glycosyltransferase and Signaling Functions

GO enrichment analysis identified six significantly enriched molecular function terms among *JAG1* candidate targets (FDR < 0.05; Figure 3F; Table S9, S10). The most significantly enriched was ADP binding (GO:0043531; 39 genes, 3.86-fold enrichment, FDR = 6.85 x 10^-11^) representing NB-ARC domain-containing proteins involved in intracellular signaling. UDP-glycosyltransferase activity (GO:0008194; 24 genes, 3.30-fold enrichment, FDR = 2.63 x 10^-5^) and protein dimerization activity (GO:0046983; 39 genes, 2.11-fold enrichment, FDR = 6.32 x 10^-4^) were also enriched, along with monooxygenase activity (GO:0004497; 23 genes, 2.23-fold enrichment, FDR = 0.014), iron ion binding (GO:0005506; 28 genes, 1.92-fold enrichment, FDR = 0.032) and oxidoreductase activity acting on paired donors (GO:0016705; 23 genes, 1.99-fold enrichment, FDR = 0.045). No biological process or cellular component terms reached significance.

### Auxin and Salicylic Acid Pathways Were Enriched Among *JAG1* Candidate Targets

Mapping KEGG Ortholog identifiers to soybean genes identified 459 unique hormone-related genes across eight pathways (Table S12). When all hormone genes were analyzed together, there was no significant enrichment among *JAG1* candidate targets (fold = 1.34, P = 0.054). However, testing each pathway individually revealed specific enrichment of auxin and salicylic acid signaling components (Figure 3G).

Auxin signaling genes showed 1.78-fold enrichment among *JAG1* candidate targets (15 observed vs 8.41 expected; P = 0.022; Figure 3G). Of these 15 auxin-related *JAG1* candidate targets, 8 (53%) were differentially expressed, all upregulated in narrow leaf genotypes. Key auxin-related *JAG1* candidate targets include *GH3* auxin-responsive conjugation enzymes (*Glyma.10G019700 and Glyma.12G103500*), *AUX1/LAX* auxin influx carriers (*Glyma.14G060700, Glyma.01G114000 and Glyma.02G255800*) and *SAUR* family proteins (*Glyma.05G196300, Glyma.06G278400 and Glyma.06G006800)* (Table S11). Direction-specific analysis revealed even stronger enrichment where auxin genes were 2.07-fold enriched among upregulated DE genes (8 observed vs 3.87 expected; P = 0.042) but showed no enrichment among downregulated genes (0.48-fold; P = 0.88) (Table S13).

Similarly, Salicylic acid (SA) signaling genes showed the strongest pathway-specific enrichment among *JAG1* candidate targets (3.99-fold; 4 observed vs 1.0 expected; P = 0.016). Two of these four SA genes were differentially expressed (*Glyma.11G183700* and *Glyma.18G020900*) and upregulated in narrow-leaved genotypes. Jasmonic acid pathway genes showed modest, but non significant enrichment (2.41-fold; P = 0.082). No other hormone pathways were signfificantly enriched (all P > 0.4; Table S14).

### *JAG1* Repression Machinery Was Available at TP1 in Both Broad and Narrow Leaves

All 12 *TPR* genes were expressed in developing leaves, with 10 showing robust expression (mean log_2_CPM > 3) at TP1 (Figure 4A; Table S7). The primary TOPLESS orthologs *GmTPR1* (*Glyma.08G214600*) and *GmTPR2* (*Glyma.07G028000*) showed the highest expression among *TPR* family members (mean log_2_CPM = 7.0 and 7.2, respectively). Five *TPR* genes showed significantly higher expression in narrow-leaved genotypes at TP1 (FDR < 0.05) including both *GmTPR1* and *GmTPR2* (Table S8).

**Figure 4.**
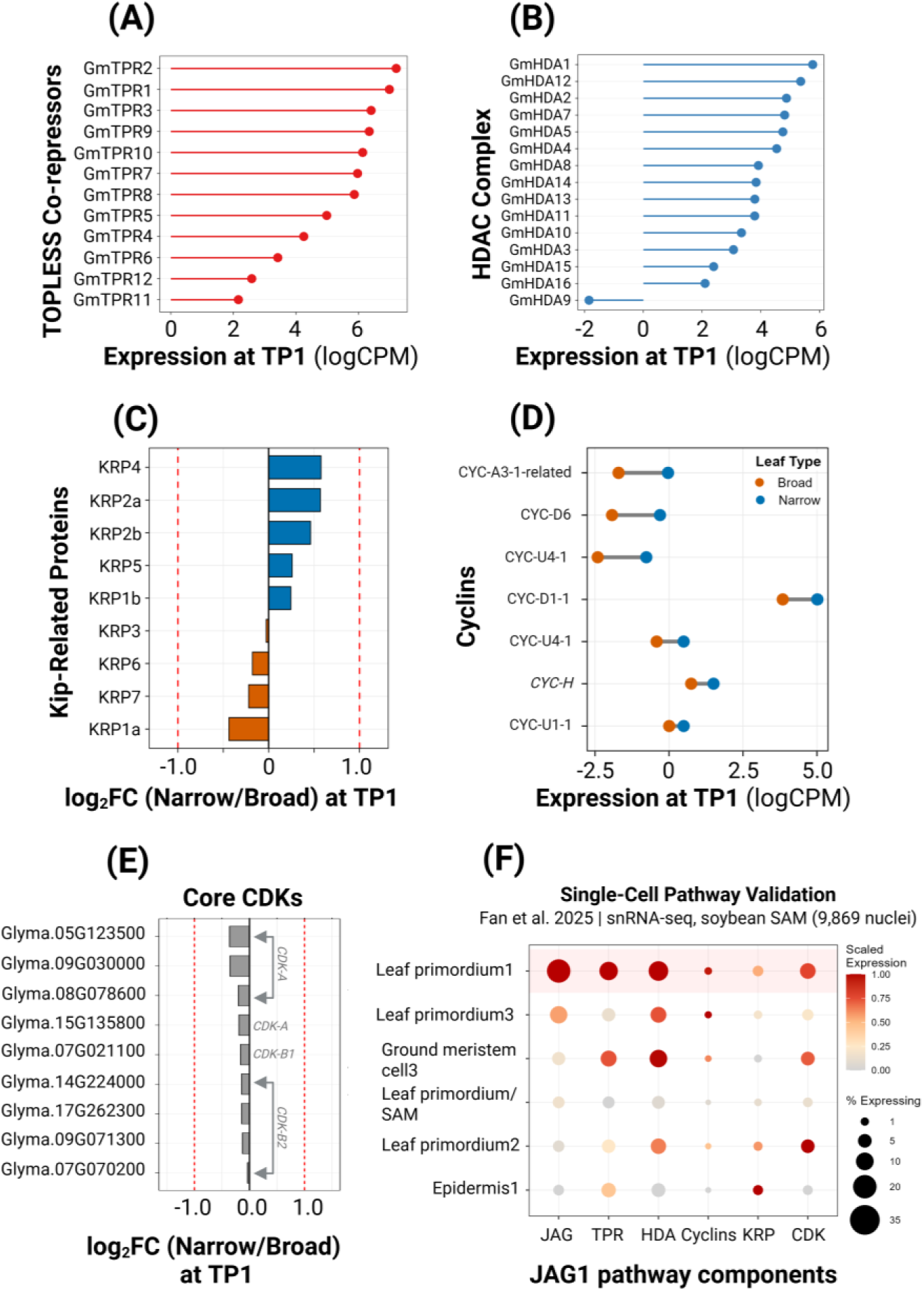
Functional analysis of GmJAG1 candidate targets and cell cycle regulation. **(A)** Expression of TOPLESS-Related Protein (TPR) co-repressor genes at TP1. Lollipop plot shows log_2_CPM expression levels. **(B)** Expression of HDAC complex genes at TP1. Lollipop plot shows log₂CPM expression levels of all 15 HDAC complex genes. **(C)** KRP (Kip-Related Protein) gene fold changes at TP1. Bar plot shows log₂ fold change (Narrow/Broad) for all 9 KRP genes. **(D)** Cyclin gene expression comparison between leaf types at TP1. Paired dot plot shows expression (log₂CPM) of 7 JAG1-targeted cyclin genes in Broad (orange) and Narrow (blue) genotypes. **(E)** Core cell cycle CDK gene fold changes at TP1. Bar plot shows log₂ fold change (Narrow/Broad) for 9 core CDKs that partner with JAG1-targeted Cyclins. Red dashed lines indicate |log₂FC| > 1 significance threshold. **(F)** Seurat-style dot plot showing mean expression of six pathway component families across selected SAM cell-type clusters from published snRNA-seq data (Fan et al. 2025; 9,869 nuclei). Dot size represents mean percent of cells expressing each gene family; dot color represents scaled mean normalized expression.

Similarly, we identified 15 HDAC complex genes in soybean, comprising nine catalytic subunits (orthologs of Arabidopsis RPD3/HDA6, HDA15, HDAC1 and HDAC11) and six regulatory subunits (SAP18 and SAP30L) (Table S7). At TP1, 14 of 15 HDAC complex genes showed detectable expression (log_2_CPM > 0) with expression levels ranging from 2.1 to 5.8 log_2_CPM (Figure 4B). The remaining gene (*GmHDA9*, a SAP18 ortholog) showed minimal expression at TP1 but increased at later timepoints.

### *KRP* Cell Cycle Inhibitors Were Not Regulated Differently in Broad and Narrow Leaves

Using two independent methods (see Materials and Methods section), we identified nine *KRP* genes in soybean (*GmKRP1a* through *GmKRP7;* Table S4).

All nine genes were expressed in leaf tissue with mean CPM values ranging from 2.1 (*GmKRP7*) to 22.3 (*GmKRP1b*) (Table S5). Cross-referencing with published binding data showed that eight of nine *KRP* genes (88.9%) had evidence of *GmJAG1* binding (Table S4). Despite this binding evidence, none of the nine *KRP* genes met our criteria for differential expression between broad and narrow leaf genotypes in pairwise comparisons. At TP1 when *GmJAG1* is maximally expressed, none of the eight *KRP*s with binding evidence showed significant differences (all FDR > 0.05), with the largest |log_2_FC| being only 0.58 (*GmKRP4*). Only *GmKRP2a* which lacked binding evidence showed a marginally significant difference at TP1 (FDR = 0.027) (Figure 4C; Table S6).

### Cyclins were *JAG1* Targets Up-regulated in Narrow Leaves

In contrast to the KRPs, we found that soybean cyclins are regulated by *GmJAG1*. 101 cyclins were identified (40 D-type, 18 B-type, 15 A-type, 9 U-type, 5 T-type, 2 H-type and 12 other/unknown) of which 94 were expressed in our dataset (Table S15). Among those, seven were identified as *JAG1* candidate targets with binding evidence which were two D-type cyclins (*CYCLIN-D1-1*, *Glyma.08G291000*; *CYCD6*, *Glyma.13G247600*), two *CYCLIN-U4-1* paralogs (*Glyma.03G107800*, *Glyma.07G119100*), *CYCLIN-A3-1* (*Glyma.06G044000*), *CYCLIN-H* (*Glyma.17G201300*) and *CYCLIN-U1-1* (*Glyma.13G306600*) (Table S16). All were upregulated in narrow-leaved genotypes. *CYCLIN-A3-1* showed the strongest fold-change (log_2_FC = 1.74) followed by *CYCD6* (log_2_FC = 1.65), *CYCLIN-U4-1* paralogs (log_2_FC = 0.94-1.50), *CYCLIN-D1-1* (log_2_FC = 1.16), *CYCLIN-H* (log_2_FC = 0.68) and *CYCLIN-U1-1* (mean log_2_FC = 0.60; maximum log_2_FC = 2.05 in PI547745 vs PI532462A) (Figure 4D).

Similarly, we identified 46 CDK genes in soybean, of which 40 were expressed in our dataset (Table S17). Core cell cycle CDKs were identified based on Arabidopsis ortholog mapping yielding 13 genes. Among them, 7 were *CDKA*, 1 *CDKB1* and 5 *CDKB2* orthologs. Of these, 9 were expressed in developing leaves (4/7 *CDKA*, 1/1 *CDKB1* and 4/5 *CDKB2*). None of the 9 expressed core CDKs were differentially expressed between narrow and broad leaf genotypes, with maximum |log_2_FC| of 0.36 at TP1 (Figure 4E, Table S17).

### Single-Cell Transcriptomics Validated Spatial Co-localization of the *JAG1* Repression Pathway

To determine whether the *JAG1* repression pathway components identified from bulk RNA-seq co-localize at the cellular level, we mapped their expression onto published snRNA-seq data from the soybean shoot apex. The dataset comprised 9,869 SAM nuclei across 16 annotated cell-type clusters and 10,084 leaf nuclei, generated from the ZH13 reference genome. Gene identifier conversion from Wm82 to ZH13 achieved 81.3% mapping for the 1,567 *JAG1* candidate targets (1,274 with ZH13 identifiers, 1,128 detected in the SAM expression matrix). *GmJAG1* was predominantly expressed in Leaf Primordium 1 (LP1), the initiating leaf primordium cluster containing 515 cells (35.1% of cells expressing, 6.8-fold enrichment over other clusters; Figure 4F, Figure S4A). The *GmJAG2* paralog showed a similar pattern (4.7% in LP1, ranked first among all clusters) but at lower absolute levels.

The full repression machinery was co-localized with *JAG1* in LP1. The 12 *TPR* co-repressor genes showed 2.1-fold enrichment in LP1 (mean 11.1% expressing vs. 5.3% in other clusters), with 10 of 12 TPRs showing peak expression in LP1 (Figure S4B). Similarly, 11 detectable HDAC complex genes showed 2.0-fold LP1 enrichment (mean 13.0% vs. 6.4%), with 8 of 11 peaking in LP1. The top expressors were *GmTPR2* (41.8% in LP1), *GmHDA3* (36.5%) and *GmHDA14* (28.9%). All three pathway components - JAG1, TPR and HDAC complex - ranked first among 16 clusters (Table S30).

In contrast, the nine *KRP* cell cycle inhibitors showed no LP1 concentration (Figure S4C). KRPs were broadly distributed across all 16 SAM clusters (mean 2.4% in LP1 vs. 1.3% in other clusters), with the highest-expressing *KRP* (*GmKRP7*, 11.8%) ranking below Epidermis1 (12.5%). Furthermore, 7 of 9 KRPs were significantly higher in mature leaf tissue than in the SAM (P < 0.05; Table S31).

The seven *JAG1*-target cyclins were expressed at very low levels in LP1 (0.0-3.1% of cells, Figure S4D). CDKs, as expected for constitutive cell cycle machinery, were more broadly expressed, with *CDKB1* (23.3%) and CDKB2-4 (16.5%) elevated in LP1.

Mapping of all 1,128 detectable *JAG1* candidate targets confirmed co-localization, with targets showing 1.97-fold higher expression in LP1 compared to other clusters (mean 3.28% vs 1.67% expressing). This expression was consistent across confidence tiers: Tier 1 (2.51-fold), Tier 2 (1.58-fold) and Tier 3 (1.97-fold) (Table S32).

### WGCNA Identified *JAG1* Target-Enriched Modules Correlated with Leaf Shape

We next used WGCNA to examine how *JAG1* candidate targets cluster within the broader transcriptional landscape. WGCNA identified 21 co-expression modules representing distinct transcriptional programs active during leaf development (Figure 5A). A total of 28,905 genes passing variance filtering were included in network construction, with soft-thresholding power of 5 achieving scale-free topology (R^2^ = 0.81, mean connectivity = 832).

**Figure 5.**
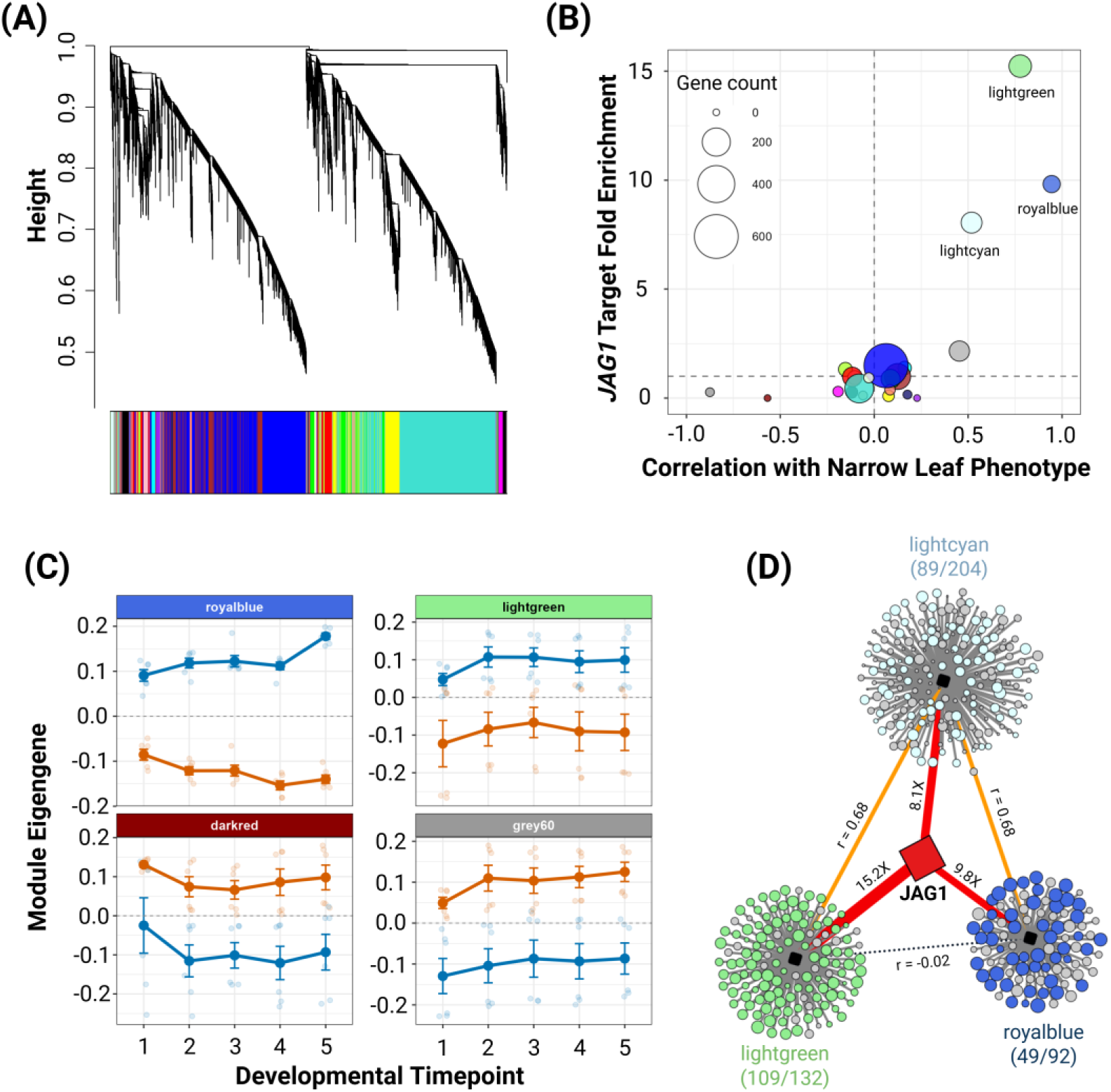
Weighted gene co-expression network analysis (WGCNA) of the soybean leaf transcriptome. **(A)** Gene dendrogram showing hierarchical clustering of 28,905 genes based on expression similarity. Color bar below the dendrogram indicates module assignments. **(B)** Scatter plot showing the relationship between module-phenotype correlation (x-axis: correlation with L:W ratio) and JAG1 candidate target enrichment (y-axis: fold enrichment). Point size represents module size; color indicates correlation direction. **(C)** Module eigengene expression trajectories across developmental timepoints (TP1-TP5) for the four modules with strongest phenotype correlations. **(D)** GmJAG1 regulatory network visualization showing target gene distribution across three highly enriched co-expression modules. Within each module, circles represent genes where colored circles are JAG1 candidate targets and gray circles are non-target genes. Node size reflects module membership strength (kME). Red arrows from GmJAG1 indicate regulatory relationships with edge width proportional to fold enrichment. Dashed lines between modules show expression correlations (orange: positive correlation).

Module-trait correlation analysis revealed that the royalblue module (92 genes) showed the strongest positive correlation with leaf L:W ratio (r = 0.94, p = 9.77 x 10^-30^). *JAG1* itself was assigned to the green module, while its targets showed strongest enrichment in distinct modules. For instance, lightgreen had a 15.23-fold enrichment (109 of 132 genes being *JAG1* candidate targets), royalblue had 9.82-fold enrichment followed by lightcyan with 8.05-fold enrichment (Figure 5B; Table S20, S22). Module eigengene trajectories confirmed that expression differences between leaf types are maintained throughout development for the most strongly phenotype-correlated modules (Figure 5C). Network visualization showed that *JAG1* candidate targets cluster within these three highly enriched modules, with regulatory connections reflecting fold enrichment (Figure 5D, Table S21).

### Multi-Evidence Integration Identified 79 High-Confidence Direct Targets

To distinguish high-confidence direct targets from indirect effects, we integrated differential expression with ChIP-seq and DAP-seq binding data, expression-phenotype correlation and WGCNA module membership. Analysis of ChIP-seq binding peak distributions showed that 52% of peaks occurred in gene body regions rather than promoters (Figure 6A). We therefore classified binding evidence based on peak location across promoter (-3000 bp to TSS), genic and downstream regions. 28,327 genes from ChIP-seq and 2,297 genes from DAP-seq identified 914 *JAG1* candidate targets (58.3%) with binding evidence. Of the 1,567 candidate targets, 885 genes (56.5%) showed ChIP-seq binding, 64 genes (4.1%) showed DAP-seq binding and 35 genes (2.2%) showed binding in both datasets (Figure 6B; Table S24-26). Correlation analysis identified 252 of 1,567 candidate targets (16.1%) whose expression significantly correlated with leaf L:W ratio (|r| > 0.5, FDR < 0.05; Figure 6C, Table S19, S24) of which 111 genes (7.1% of 1,567) also had binding evidence. Applying sequential filters for binding evidence, significant phenotype correlation and membership in WGCNA phenotype-correlated modules yielded 79 high-confidence targets supported by all four types of evidence (5.0% of 1,567; Figure 6D; Table S27, S29), including 8 assigned to the WGCNA unassigned module. These 79 genes include 15 Tier 1, 10 Tier 2 and 54 Tier 3 genes, with 4 supported by both ChIP-seq and DAP-seq binding evidence.

**Figure 6.**
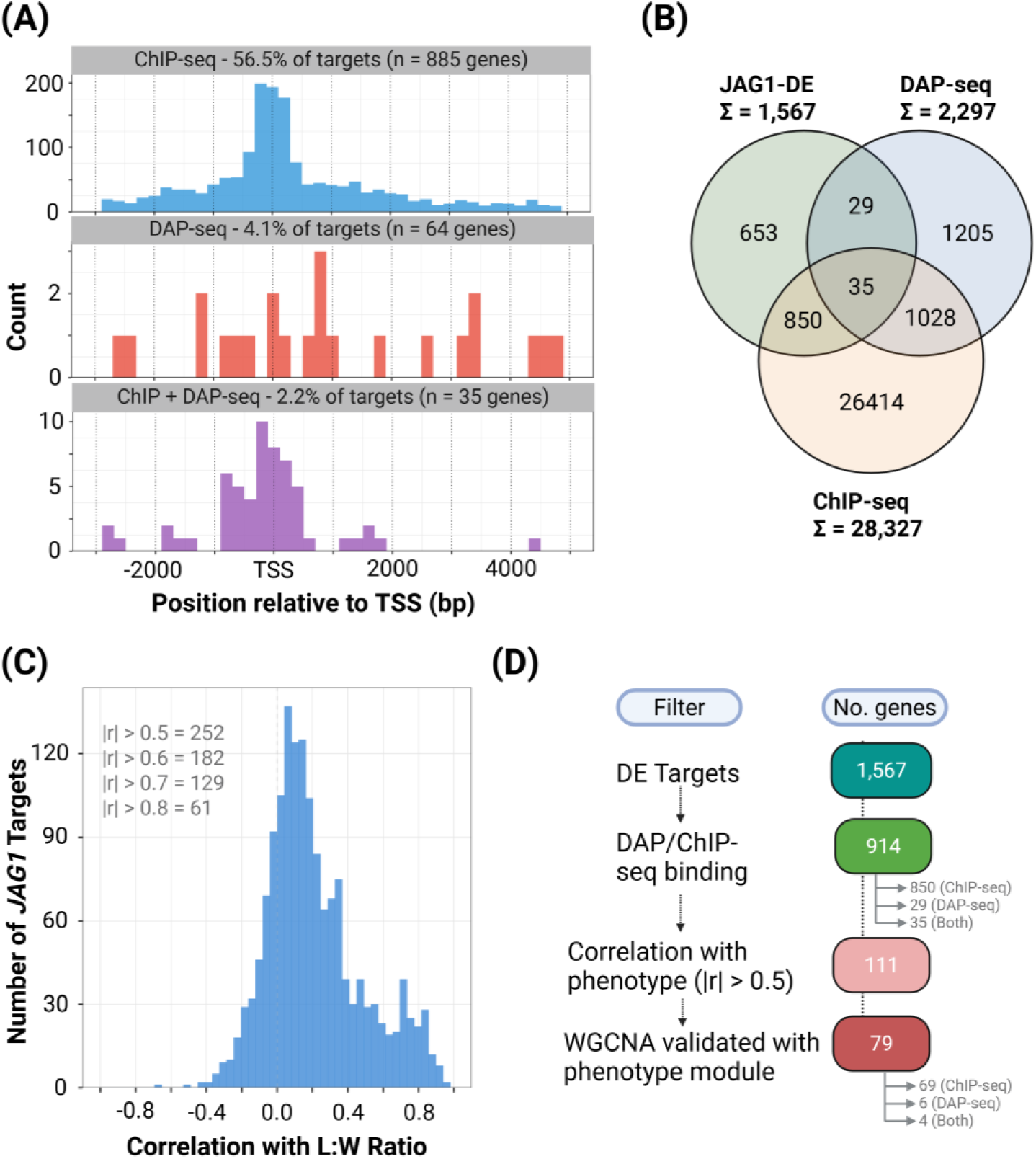
Multi-evidence validation of GmJAG1 target genes. (A) Positional distribution of GmJAG1 binding peaks relative to transcription start site (TSS). Histograms show the distribution of ChIP-seq binding peaks (Huang et al., 2021), DAP-seq binding peaks (Wang et al., 2024) and genes with both ChIP-seq and DAP-seq binding evidence (highest confidence; bottom, purple) across a window from -3000 bp (upstream/promoter) to +5000 bp (gene body/downstream). (B) Three-way Venn diagram showing overlap between JAG1 differentially expressed targets. Numbers within each region indicate gene counts. (C) Distribution of correlation coefficients between JAG1 target gene expression and leaf length-to-width (L:W) ratio. (D) Evidence integration flow diagram showing sequential filtering of JAG1 candidate targets through multiple validation layers.

Functional characterization based on Arabidopsis ortholog annotation showed that the 79 targets span varying biological processes (Figure 7; Table S29). The largest category was metabolic enzymes (15 genes, 19%) including glycosyltransferases, cytochrome P450s and oxidoreductases. Hormone signaling genes (9 genes, 11%) included components of auxin (*NPH3* and *CYP83B1*), jasmonate (*JAZ1*), cytokinin, gibberellin, ABA and brassinosteroid pathways. Defense/NBS-LRR genes (7 genes, 9%), receptor kinases (5 genes, 6%) and transcription/chromatin regulators (7 genes, 9%) were also well represented. Temporal expression profiles showed that most targets were upregulated across all five developmental timepoints (TP1-TP5) with the strongest and most consistent changes at TP1 corresponding to peak *GmJAG1* expression. One exception was *PhyE2* (*Glyma.15G196500*; log_2_FC = -2.4 at TP1), which appeared downregulated in the pooled comparison driven by near-absent expression in PI 612713B while PI 547745 expressed at levels comparable to broad genotypes.

**Figure 7.**
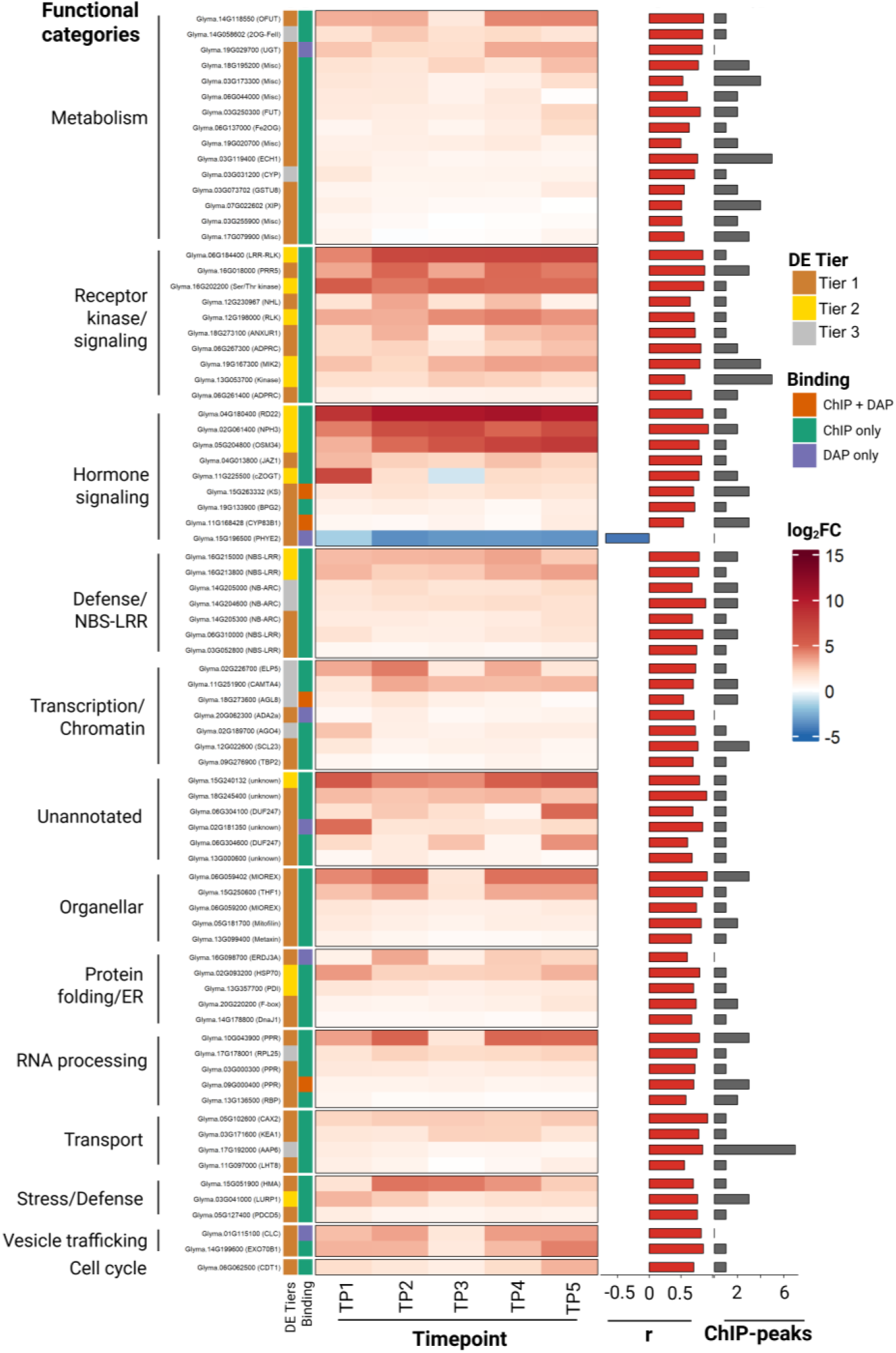
Functional characterization of 79 high-confidence GmJAG1 target genes. Heatmap showing log_2_ fold change (log_2_FC) of 79 validated GmJAG1 targets across five developmental timepoints (TP1-TP5) comparing narrow leaf versus broad leaf soybean genotypes. Genes are grouped by functional category based on Arabidopsis ortholog annotation. Left annotations indicate differential expression tier and binding evidence class. Right annotations show Pearson correlation coefficient with leaf length-to-width ratio (r) and number of ChIP-seq binding peaks. Red-blue color scale represents upregulation and downregulation, respectively. TP1 corresponds to the developmental stage of peak GmJAG1 expression.

Among the most strongly induced targets were *RD22* (*Glyma.04G180400*; log_2_FC = 12.2 at TP1), an ABA-responsive dehydration gene, and trans-zeatin O-glucosyltransferase (*Glyma.11G225500*; log_2_FC = 10.1 at TP1), a cytokinin-inactivating enzyme. In the single-cell transcriptomic data, 43 of the 79 targets were detectable in the SAM expression matrix, with 31 (72%) expressed in LP1 at low levels (mean 2.69%).

### qRT-PCR Independently Validated Key RNA-seq Findings

To confirm the key RNA-seq findings by an independent method, we performed qRT-PCR on eight target genes across all four genotypes at TP1 (Table S34; Figure S6). qRT-PCR confirmed that neither *GmJAG1* nor *GmJAG2* differed between leaf types (both P >0.05; Figure 3A). Among D-type cyclins, *CYCLIN-D1-1*, the most abundantly expressed *JAG1*-target cyclin (51.8 CPM), showed significant upregulation in narrow-leaf genotypes (2.0-fold, P = 0.018), while *CYCD6* trended in the expected direction (1.2-fold, P = 0.088), consistent with its low expression level (<1 CPM). Both KRP negative controls showed no differential expression (*KRP1b* P = 0.30; *KRP4* P = 0.46), independently confirming the absence of KRP regulation. RD22 was massively upregulated in narrow genotypes (>3,000-fold, P < 0.001), while NPH3 showed genotype-specific rather than leaf-type-dependent expression, being undetectable in PI 532462A but expressed at comparable levels in the other three genotypes. Overall, qPCR fold changes correlated strongly with RNA-seq estimates (Pearson r = 0.99, P < 0.001, slope = 1.07; Figure S6).

## Discussion

A key challenge in plant biology is understanding how transcription factors translate genetic variation into differences in plant form. Genome-wide binding studies can show where transcription factors interact with DNA, but distinguishing functional targets from non-productive binding events requires independent approaches (Higgins et al., 2025). Here, we used two complementary strategies to dissect the regulatory role of *GmJAG1* in soybean leaf development. First, we took a hypothesis-driven approach, examining the cell cycle genes and repression machinery that mediate *JAG* function in Arabidopsis including *KRP* inhibitors, cyclins, *CDK*s, *TOPLESS* co-repressors and *HDA*s. We tested whether the established regulatory mechanism is conserved in soybean. Second, we employed a data-driven approach, using comparative transcriptomics between broad and narrow-leaved genotypes to let natural variation at the *GmJAG1* locus reveal its functional targets. These two approaches show that *GmJAG1* does not regulate *KRP*s despite evidence of promoter binding but instead may control D-type cyclins and that it regulates growth beyond cell cycle control including auxin signaling, salicylic acid pathways and diverse developmental regulators.

### Natural Variation as a System for Candidate Target Identification

Previous studies of *GmJAG1* used ChIP-seq and DAP-seq to map genome-wide binding sites (Huang et al., 2021; Wang et al., 2024). While these approaches identify where *GmJAG1* binds DNA, they cannot tell us which binding events lead to changes in gene expression. The D9H mutation resides in the EAR repression motif (Jeong et al., 2012) that recruits the TOPLESS co-repressor complex for transcriptional silencing (Szemenyei et al., 2008), not in the C2H2 zinc finger DNA-binding domain. Both protein variants should therefore recognize and bind the same genomic targets but possibly differ in co-repressor recruitment. De-repression of target genes in narrow genotypes, consistent with our finding that most differentially expressed genes were upregulated (65.7% of 1,098). The remaining downregulated genes likely represent indirect effects, as the upregulated proportion increased with evidence stringency: 78 of 79 high-confidence targets (98.7%) were upregulated, with only *PhyE2* showing apparent downregulation (see results). Our approach exploits this natural variation to find genes whose expression changes when *JAG1* repression function is lost. Genes showing both binding evidence and differential expression are therefore good candidates for direct targets.

This apex-restricted expression aligns with the strong meristem expression reported by Jeong et al. (2012), who also detected *GmJAG1* in open flowers. In Arabidopsis, *JAG* is expressed in lateral organ primordia rather than the meristem itself (Ohno et al., 2004), suggesting that meristem expression may be a derived feature in soybean. Despite this transient expression, most *JAG1* candidate targets maintained differential expression throughout development (Figure 3E).

### *KRP* Cell Cycle Inhibitors Were Not Transcriptionally Regulated Despite Binding Evidence

We examined whether the Arabidopsis *JAG*-*KRP* regulatory connection (Schiessl et al., 2014) is conserved by analyzing the complete nine-member soybean *KRP* family.

The Arabidopsis *JAG-KRP* regulatory connection provided a clear prediction. If conserved in soybean, *KRP* genes should be de-repressed in narrow-leaf genotypes lacking functional *JAG1*. None showed that pattern (Figure 4C) despite binding evidence (8 of 9 KRPs) and available repression machinery. Even in Arabidopsis, this prediction was only partially met where *KRP2* showed de-repression in *jag* mutants, but *KRP4* did not, despite direct binding and gain-of-function repression evidence (Schiessl et al., 2014).

A potential explanation is functional redundancy with the *JAG2* paralog, which shares ∼91% amino acid identity with *JAG1* (Figure S3). *JAG2* retains both an intact EAR motif and identical C2H2 zinc finger domain and is expressed at TP1 at similar levels in both genotypes raising the possibility that it maintains *KRP* repression when *JAG1* function is compromised. However, promoter analysis reveals no obvious basis for selective compensation where both *KRP*s and cyclins showed similar M0 motif prevalence (67% each) and binding evidence, yet only cyclins responded transcriptionally. Furthermore, snRNA-seq data from the soybean shoot apex (Fan et al., 2025) showed no *KRP* concentration in LP1, the cluster where *JAG1* and its repression machinery co-localize (Figure S4), ruling out bulk tissue-averaging as an explanation and suggesting genuine regulatory divergence. Recent work on transcription factor paralogs demonstrates that proteins with nearly identical DNA-binding domains can bind the same genomic sites yet regulate largely different gene sets through ’differential usage’ of shared binding sites, mediated by as few as eight amino acid substitutions outside the binding domain (Higgins et al., 2025). *JAG1* and *JAG2* share identical C2H2 zinc finger and EAR motif sequences (Figure S3) but differ at approximately 9% of residues in flanking regions. These non-conserved residues may determine which commonly bound targets are functionally responsive to each paralog, potentially explaining why cyclins but not KRPs respond when *JAG1* repression is lost while *JAG2* remains functional. Testing this model will require JAG2-specific ChIP-seq to determine whether the two paralogs share binding sites but differ in transcriptional outcomes.

### D-type Cyclins - Transcriptional Targets of *GmJAG1*

Extending our analysis of cell cycle regulators, we examined whether *GmJAG1* controls cell cycle activators rather than inhibitors. Seven cyclins were *JAG1* candidate targets with binding evidence, including D-type cyclins *CYCD6* and *CYCLIN-D1-1*, all upregulated in narrow leaf genotypes (Figure 4D). D-type cyclins are rate-limiting for the G1-to-S phase transition and promote cell proliferation (Dewitte et al., 2003; Menges et al., 2006). Interestingly, the same D-type cyclin subfamilies targeted by *GmJAG1*- *CYCD1* and *CYCD6*- are also direct targets of the soybean *PLATZ* transcription factor, which activates their expression to promote seed size through cell proliferation (Hu et al., 2023), independently validating these cyclins as key regulatory nodes in soybean organ development. Their upregulation when *JAG1* repression is lost suggests that *GmJAG1* normally represses cyclin expression during leaf development. qRT-PCR independently confirmed D-type cyclin upregulation and KRP non-regulation. Consistent with this, all seven were expressed at very low levels in LP1 in the Fan et al., 2025 snRNA-seq dataset, which was generated from a broad-leaf genotype with functional *JAG1*, as expected under active repression by the JAG1-TPR-HDA complex co-localized in this cluster.

The upregulation of D-type cyclins in narrow leaf genotypes might seem paradoxical-elevated cell cycle activators accompanying a narrower leaf. However, leaf shape reflects the spatial organization of cell division, not simply its quantity. Loss of *JAG* function in Arabidopsis causes reversion to isotropic, meristem-like growth rather than simply reducing cell number (Schiessl et al., 2012). In soybean, narrow genotypes show 30% higher LMA (58.9 vs. 45.5 g m⁻²; P = 0.004; Figure 1E) and increased leaf thickness (Tamang et al., 2023), indicating a dorsoventral growth component. Leaf width, however, is primarily determined by oriented cell divisions at lateral margins (Horiguchi et al., 2005; Kinoshita et al., 2022), and *GmJAG1* acts exclusively at the shoot apex when these division patterns are established. De-repression of cyclins at this critical window may disrupt the coordinated program that normally promotes lateral expansion, redistributing growth from width to thickness- consistent with the absence of Leaf x Day interaction for L:W ratio (all P > 0.2). This interpretation is further supported by *CDK* analysis where none of the nine core CDKs were differentially expressed (maximum |log2FC| = 0.36), indicating that *GmJAG1* controls cell division at the level of upstream activators rather than kinases or their inhibitors (Table S18).

Together, these results reveal a divergence in regulatory logic over ∼90 million years of independent evolution (Grant et al., 2000). In Arabidopsis, *JAG* uses double-negative logic repressing inhibitors to permit division whereas in soybean, *GmJAG1* appears to directly repress activators to coordinate division patterns. Schiessl et al. (2014) identified *CYCD3;3* among *JAG*-repressed genes in Arabidopsis indicating that cyclin repression is not entirely novel but was a minor component of a predominantly *KRP*-centered mechanism. In soybean, this relationship appears inverted where cyclin repression is a primary transcriptional response while KRPs are unaffected. Phylogenetic analysis confirmed that the *CYCD1* and *CYCD6* subclades containing the two *GmJAG1*-regulated cyclins are deeply divergent from *CYCD3*, which contains *AtCYCD3;3* (Figure S5), suggesting that the relative importance of these two regulatory arms has shifted during legume evolution.

Interestingly, cyclin upregulation in narrow-leaf genotypes is not unique to soybean. In wheat, narrow-leaf mutant *nl1* caused by a mutation in the CDC48-like protein *NL1* also shows cyclin E upregulation (Li et al., 2026). That independent causal genes in a monocot and dicot converge on cyclin upregulation in narrow-leaf genotypes suggests this is a conserved downstream feature of leaf width determination.

### Auxin-SA Co-enrichment Suggests Coordinated Growth-Defense Regulation

Examination of the1,567 *JAG1* candidate target genes revealed broader regulatory patterns. Auxin and SA signaling were specifically enriched among *JAG1* candidate targets, a signal apparent only when pathways were tested individually (Figure 3G). Especially, auxin enrichment was stronger among upregulated DE genes (2.07-fold, P = 0.042) than among all targets (1.78-fold), consistent with de-repression when *JAG1* function is lost. These findings link JAG-family transcription factors to auxin pathway regulation, though we note that the unrelated Arabidopsis gene JAGGED LATERAL ORGANS (JLO), an *LBD* family member whose gain-of-function mutant produces lobed lateral organs, also affects auxin transport through regulation of *PIN* carriers (Borghi et al., 2007). The co-targeting of cyclins and auxin genes is biologically coherent where auxin promotes cell division through the *ARGOS-ANT-CYCD3* pathway (Hu et al., 2003), and *CYCD2;1* modulates auxin-induced lateral root formation (Sanz et al., 2011). De-repression of both cyclins and auxin signaling in narrow leaf genotypes may amplify effects on cell proliferation.

SA pathway genes showed the strongest pathway-specific enrichment (3.99-fold, P = 0.016; Figure 3G). Direct metabolite measurements in the wheat *nl1* mutant confirmed a 1.67-fold higher SA in narrow leaves (Li et al., 2026), providing cross-species support that SA accumulation accompanies narrow-leaf phenotypes. Beyond its role in defense, SA negatively regulates cell expansion where SA hyperactivation suppresses compensated cell enlargement in *an3* mutants (Fujikura et al., 2020), which, like our narrow leaf genotypes, show altered leaf dimensions. Auxin and SA tend to be antagonistic where SA inhibits growth partly by repressing auxin signaling (Wang et al., 2007), forming what has been described as a “seesaw” balancing growth and defense (Deng and He, 2024; Han et al., 2025). The convergence of auxin and SA enrichment with mechanistic links to cell proliferation warrants further investigation.

Beyond hormone signaling, GO enrichment of *JAG1* candidate targets identified UDP-glycosyltransferases, protein dimerization activity and monooxygenase activity as significantly enriched categories (Table S9), consistent with coordinated regulation of cell wall remodeling, protein interactions, and signaling during organ growth.

### An Integrated Model of *GmJAG1* Function

Integration of four complementary layers- differential expression, binding evidence, phenotype correlation and co-expression module membership- identified 79 high-confidence potential targets (Figure 6D; Table S27, S29). Several have Arabidopsis orthologs with characterized roles in organ development (Figure 7; Table S29). Together with the hypothesis-driven cell cycle analysis, these results point to *GmJAG1* as a coordinator of multiple processes during early leaf development (Figure 8). At the meristem, functional *GmJAG1* represses targets via its EAR motif, recruiting *TOPLESS* and histone deacetylases to silence gene expression. Three classes of targets emerged from the combined analyses.

**Figure 8.**
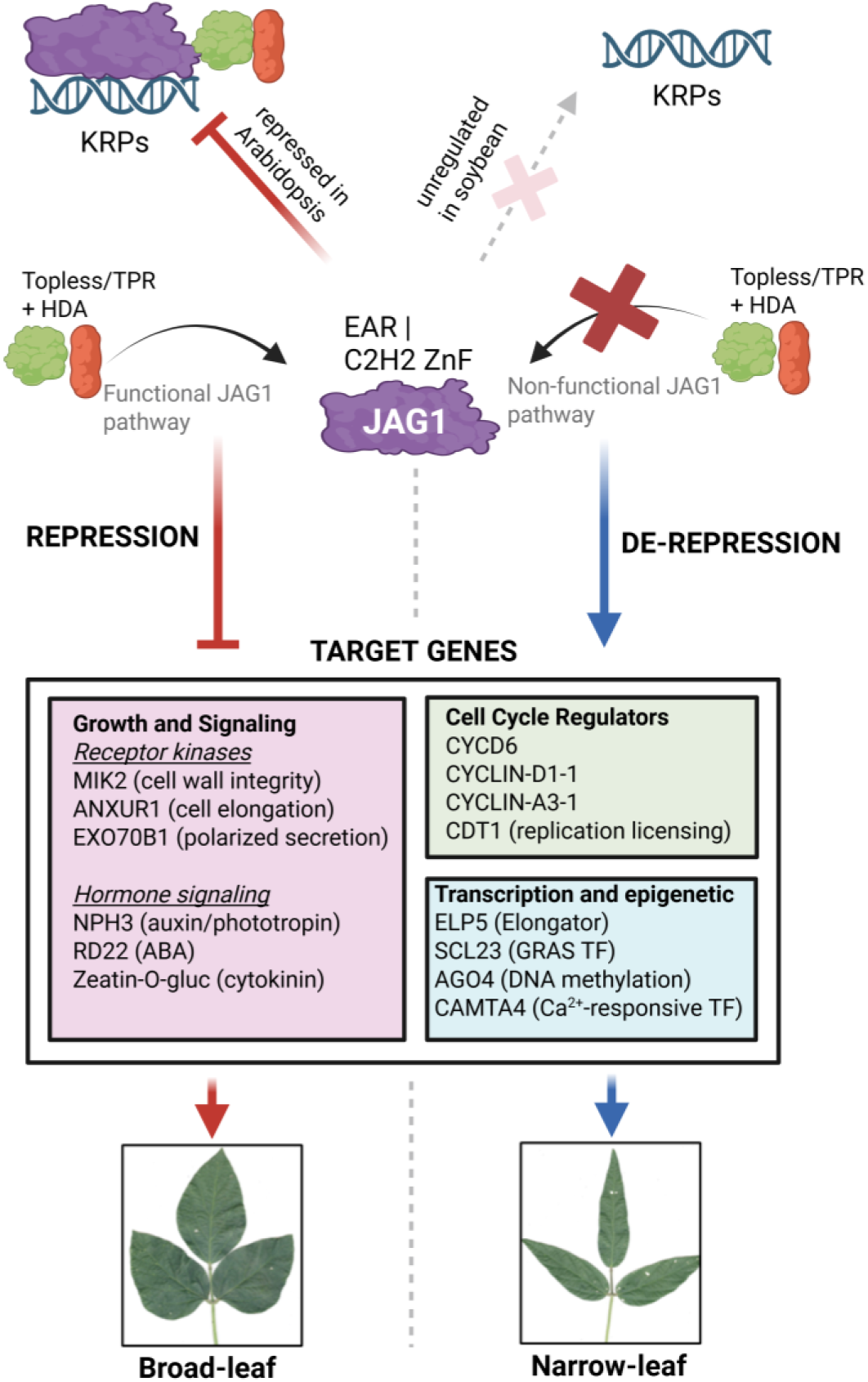
Proposed model of GmJAG1-mediated transcriptional regulation in soybean leaf development. Comparison of JAG1 regulatory function in broad leaf (left, functional JAG1) and narrow leaf (right, D9H mutation) genotypes. Blunt-ended arrow indicates active repression; red "X" indicates disrupted protein-protein interaction due to D9H mutation. Gene names represent GmJAG1 targets identified in this study.

First, cell cycle regulators: D-type cyclins and *CYCLIN-A3-1* were upregulated with binding evidence (Figure 4D), representing the primary cell cycle response to *JAG1* loss. *CDT1*, a DNA replication licensing factor, was the only cell cycle gene among the 79 high-confidence targets supported by all four evidence layers. The seven *JAG1*-targeted cyclins were supported by differential expression and binding evidence but did not meet the phenotype-correlation threshold (|r| > 0.5 with L:W ratio), consistent with their role as upstream regulators of cell division patterns rather than direct determinants of leaf shape.

Second, growth and signaling genes included receptor kinases: *MIK2* (cell wall integrity sensing; Van der Does et al., 2017), an *ANXUR1*-related receptor-like kinase (cell elongation; Guo et al., 2009), *EXO70B1* (polarized secretion; Hála et al., 2008), and hormone signaling components spanning six pathways (Figure 7; Table S29). *NPH3*, whose Arabidopsis ortholog mediates phototropin-dependent leaf flattening (Kozuka et al., 2013), showed the strongest phenotype correlation. *RD22* and a zeatin O-glucosyltransferase were the most strongly induced targets where the latter parallels *CKX*-mediated cytokinin reduction in wheat *nl1* narrow-leaf mutant (Li et al., 2026).

Third, transcription and epigenetic regulators may explain how transient *GmJAG1* expression establishes lasting developmental programs: *ELP5* (Elongator complex subunit; Nelissen et al., 2005), *SCL23* (*SHR*-*SCR* network; Cui et al., 2014; Yoon et al., 2016), AGL8 (meristem identity; Ferrándiz et al., 2000), *AGO4* (siRNA-directed methylation) and *ADA2a* (histone acetylation). De-repression of these regulators when *JAG1* function is lost could propagate altered gene expression beyond the meristem stage, consistent with the continued differential expression observed at TP2-TP5.

Taken together, cell cycle regulators control where and how cells divide, growth and signaling components provide spatial inputs that coordinate growth patterns, and transcriptional/epigenetic regulators maintain altered expression beyond the transient window of *GmJAG1* activity. *GmJAG1* thus coordinates multiple processes during early leaf development (Figure 8), consistent with the large phenotypic effect (>70% of variance) of this single locus (Tamang et al., 2026c) clarifying why the narrow leaflet allele reduces leaf area index without compromising yield (Tamang et al., 2026a).

## Study Limitations

Although we have integrated several strands of evidence to support the JAG1-mediated transcriptional model in soybean, the study has limitations. First, the available *GmJAG1* binding data comes from protoplast-based ChIP-seq and *in vitro* DAP-seq rather than ChIP-seq performed directly on developing leaf primordia, which may not fully capture tissue-specific binding contexts. Second, our transcript comparisons used diverse germplasm (plant introductions and an elite cultivar) rather than transgenic overexpression or knockout lines, so we cannot definitively establish causation.

Our bulk RNA-seq approach samples entire leaf primordia, which may dilute region-specific transcriptional signals. Leaf width is primarily determined by cell division patterns at the lateral margins; in narrow leaves, margin cells may exhibit arrested or reoriented division while the central blade continues normal development.

Because our tissue collection homogenized the primordium, we may miss margin-localized *JAG1* effects that are muted by signals from other leaf regions. This means that absence of differential expression does not definitively rule out region-specific regulation. However, single-cell transcriptomic analysis of the soybean shoot apex (Fan et al., 2025) showed no *KRP* concentration in the *JAG1-*expressing LP1 cluster, arguing against a simple dilution explanation for *KRP* non-responsiveness. Nonetheless, margin-specific regulation below the resolution of current single-cell cluster annotations cannot be fully excluded. The contrast between our negative *KRP* results and positive cyclin results is therefore particularly informative where cyclins show differential expression strong enough to emerge from bulk tissue averaging, whereas any *KRP* regulation, if present, must be either absent or too spatially restricted to detect.

Finally, the experimental design involved confounding between genetic line and leaf type (each line exhibits only one phenotype), though consistency of results across four independent pairwise comparisons mitigates this concern.

## Future Directions

Functional validation of the top-ranked targets (Table S28) with characterized Arabidopsis homologs- *NPH3* (phototropin-mediated leaf flattening), *MIK2* (cell wall integrity sensing), *RD22* (ABA-mediated stress signaling) and *SCL23* (GRAS transcription factor involved in bundle sheath development and leaf size regulation)- through CRISPR-mediated knockout or overexpression would test whether these genes contribute to the narrow leaf phenotype. *NPH3* is a strong candidate given the strongest phenotype correlation (r = 0.93) and its established role in leaf blade flattening. Spatial transcriptomics or laser capture microdissection of leaf margins versus central blade would resolve whether *JAG1* effects are spatially restricted, potentially revealing regulatory effects not detectable in bulk tissue. ChIP-seq of *JAG2* to determine its binding specificity would test whether *JAG2* compensation explains the absence of *KRP* regulation. Recently registered near-isogenic lines sharing ∼95% genetic similarity with the recurrent parent and differing principally at the *GmJAG1* locus (Tamang et al., 2026b) would enable definitive validation of these targets by eliminating background genetic variation as a confounding factor. Finally, the defined molecular targets of *GmJAG1* may inform precision breeding strategies for canopy optimization, as introgression of the narrow leaflet allele reduces leaf area index by ∼13% without yield penalty under field conditions (Tamang et al., 2026a).

## Conclusion

Using natural leaf shape variation, we identified 1,567 *GmJAG*1 candidate targets and found that seven cyclins including the G1/S regulatory cyclins, but not *KRP* inhibitors, show differential expression consistent with *JAG1*-mediated regulation despite binding evidence at both classes of genes. This pattern suggests that the regulatory logic connecting *JAG* transcription factors to cell cycle control may differ between soybean and Arabidopsis, though functional redundancy with the *JAG2* paralog cannot yet be excluded. Cyclins, auxin signaling and salicylic acid pathway genes are co-enriched among targets, suggesting that *GmJAG1* coordinates cell cycle and hormone signaling during early leaf development. Multi-evidence filtering identified 79 high-confidence direct targets spanning hormone signaling, receptor kinases, defense and transcriptional regulation. These candidates are priorities for functional validation, particularly in near-isogenic backgrounds, and could inform breeding strategies for soybean canopy optimization.

## Materials and Methods

### Plant Materials and Experimental Design

Four soybean (*Glycine max*) genotypes with contrasting leaf morphology were selected for this study based on their leaf length-to-width (L:W) ratios: two narrow-leaf genotypes, PI 547745 (L:W = 2.16) and PI 612713B (L:W = 2.36) and two broad-leaf genotypes, LD11-2170 (L:W = 1.59) and PI 532462A (L:W = 1.55). Plants were grown in controlled environment chambers (Conviron PGR15, Conviron, Winnipeg, BM, Canada) under a 14-hour photoperiod at 30^°^ C day/20^°^ C night temperatures. Leaf tissue was harvested at five developmental timepoints (TP1-TP5) spanning early leaf primordium initiation through full expansion of the second trifoliate leaf. Three biological replicates were collected for each genotype-timepoint combination, resulting in 60 samples in total. These samples were collected across two batches where two genotypes (PI 547745 and LD11-2170) were sampled at TP2-TP5 (24 samples) in batch 1. In batch 2 (36 samples), TP1 was added for these genotypes (6 additional samples) and two additional genotypes (PI 612713B and PI 532462A) were sampled at all five timepoints TP1-TP5 (30 samples).

The same four genotypes were also grown under field conditions at the SoyFACE experimental farm, University of Illinois at Urbana-Champaign, during the 2022 growing season. Four-row plots (2.4 m, 0.76 m row spacing) of each genotype were arranged in a randomized complete block design (RCBD) with three replicate blocks.

### Leaf Morphological Measurements

On the chamber-grown samples, leaf length and width of the unifoliate leaf and middle trifoliate leaflet of the first and second leaves were manually measured from day 1 through day 16 after leaf emergence (n = 10 plants per genotype). Length-to-width (L:W) ratio was calculated for each measurement. Differences between broad and narrow leaf types were tested using two-way fixed-effects with leaf type (broad vs. narrow) and day after emergence (as a categorical factor) as main effects, including the leaf type x day interaction term, fitted separately for each leaf node and trait using the *aov* function in R.

Leaf thickness was measured from cross-sections prepared as described in Tamang et al. (2023), at 10 positions per biological replicate (5 on each side of the midrib; n = 3 per genotype).

In field grown plants, leaf area, length, and maximum width were measured using an LI-3000C portable leaf area meter (LI-COR Biosciences, Lincoln, NE, USA). Leaf mass per unit area (LMA) was determined as leaf dry weight (after 7 days drying at 65^°^ C) divided by fresh leaf area. Differences between broad and narrow leaf types were tested using RCBD ANOVA on plot-level means. The model included leaf type (broad vs. narrow) as a fixed factor and block (n = 3) as a blocking factor, fitted using the *aov* function in R.

### *GmJAG1* Sequencing and qRT-PCR Validation of RNA-Seq Findings

To confirm the *GmJAG1* allele in each genotype, the *GmJAG1* coding region was amplified by PCR from genomic DNA using a forward primer (5’-AAAAGCTTTAGTTTTATCCCTACC-3’) and a reverse primer (5’-CGTGTCTTTATGATGATACCAC-3’). PCR products were sequenced by Sanger sequencing at the Roy J. Carver Biotechnology Center, University of Illinois.

To validate key RNA-seq findings by an independent method, qRT-PCR was performed on eight genes across all four genotypes at TP1 (three biological replicates each). Target genes were selected to represent the major findings: *GmJAG1* and *GmJAG2* (paralog expression), *CYCD1-1* and *CYCD6* (D-type cyclin regulation), *KRP1b* and *KRP4* (negative controls), and *RD22* and *NPH3* (high-confidence targets). cDNA was synthesized from total RNA using the QuantiTect Reverse Transcription Kit (Qiagen). qRT-PCR was performed on a CFX Opus 96 Real-Time PCR Detection System (Bio-Rad) using SsoAdvanced Universal SYBR Green Supermix (Bio-Rad). Expression was normalized to the geometric mean of two validated reference genes (Vandesompele et al., 2002), *Actin 11* (*Glyma.18G290800*; Machado et al., 2020) and *RPL30* (*Glyma.13G318800*; Tamang et al., 2021), following MIQE guidelines. Primer sequences, amplicon sizes and amplification efficiencies are listed in Table S33. For target-reference gene pairs with efficiency differences less than 5%, the 2^(-delta delta Ct) method was applied (Livak and Schmittgen, 2001). For pairs differing by more than 5%, the Pfaffl correction was used (Pfaffl, 2001). Each biological replicate was measured with 3-5 technical replicates. Melt curve analysis confirmed single-product amplification for all primer pairs. Differences between broad and narrow leaf types were tested using two-sample t-tests on delta Ct values (Table S34).

### RNA Extraction and Sequencing

Total RNA was extracted from frozen leaf tissue using the RNeasy Plant Mini Kit (Qiagen) and assessed on a 1% agarose gel. RNA samples were submitted to the Roy J. Carver Biotechnology Center, University of Illinois, where DNase treatment was performed to remove genomic DNA contamination, and RNA integrity was verified using an Agilent Bioanalyzer. Strand-specific mRNA libraries were prepared using poly-A selection with the TruSeq Stranded mRNAseq Sample Preparation kit (Illumina; batch 1 samples) or the KAPA Hyper Stranded mRNA library kit (Roche; batch 2 samples). Libraries were pooled, quantified by qPCR and sequenced on an Illumina NovaSeq 6000 using an SP flow cell, generating 100-bp single-end reads.

### Read Processing and Quantification

Raw sequencing reads were quantified using Salmon v1.10.0 (Patro et al., 2017) with selective alignment against the soybean reference transcriptome. A decoy-aware index was constructed by combining the *Glycine max* Williams 82 transcriptome (Wm82.a6.v1) with the genome assembly (Gmax_880_v6.0) (Espina et al., 2024) obtained from Phytozome (Goodstein et al., 2012). Chromosome sequences served as decoys to improve mapping specificity and reduce spurious alignments. Quantification was performed with automatic library type detection, GC bias correction and sequence-specific bias correction. Transcript-level abundance estimates were imported into R and summarized to gene-level counts using tximport (Soneson et al., 2015) with the “lengthScaledTPM” method. Mapping rates and quality metrics were aggregated using MultiQC (Ewels et al., 2016).

### Data Processing, Quality Control and Batch Correction

Gene expression counts were normalized using the trimmed mean of M-values (TMM) method implemented in edgeR (Robinson et al., 2010). Genes with low expression were filtered using a threshold of counts per million (CPM) > 1 in at least 3 samples (Chen et al., 2016).

To validate RNA-seq data quality, we assessed expression stability of 452 established soybean housekeeping genes obtained from Machado et al. (2020), who identified these genes through meta-analysis of 1,298 publicly available RNA-seq samples. Expression stability was quantified using the coefficient of variation (CV) calculated across all 60 samples.

Samples were sequenced across two batches (batch 1 with 24 samples and batch 2 with 36 samples), with batch partially confounded with genotype and timepoint (see above). Therefore, batch effects were corrected using ComBat-seq (Zhang et al., 2020). Timepoint and leaf type were included as covariates to preserve biological signal during correction. The effectiveness of batch correction was assessed through: Principal Variance Component Analysis (PVCA) which partitions total variance into biological (timepoint, leaf type) and technical (batch) components, PCA to visualize sample clustering before and after correction, Relative Log Expression (RLE) plots to assess normalization consistency and per-gene R^2^ calculations to quantify the proportion of variance explained by batch.

### Differential Expression Analysis and *JAG1* Target Identification

Differential expression analysis was performed in R using the edgeR quasi-likelihood (QL) framework (v4.6.3; Chen et al., 2016). Gene-wise dispersions were estimated with robust empirical Bayes shrinkage using estimateDisp, and a generalized linear model was fitted using glmQLFit with robust = TRUE to moderate the quasi-likelihood dispersions toward a common value, reducing the impact of outlier genes. For each pairwise contrast (narrow vs. broad at each timepoint) and the pooled comparison, quasi-likelihood F-tests were performed using glmQLFTest. Genes with FDR < 0.05 and |log2FC| > 1 were considered significantly differentially expressed.

A tiered approach was used to identify high-confidence *GmJAG1* candidate targets based on reproducibility across independent pairwise comparisons. Three tiers were defined where Tier 1 genes were differentially expressed in 4/4 pairwise comparisons, Tier 2 genes where genes were differentially expressed in 3/4 pairwise comparisons and Tier 3 genes where genes were either differentially expressed in 2/4 pairwise comparisons or in pooled comparisons.

### Integration with Published *GmJAG1* Binding Data

To identify direct *GmJAG1* targets, we integrated our differential expression data with published binding datasets. *GmJAG1* binding data were obtained from two studies. First, Huang et al. (2021) performed ChIPmentation on soybean protoplasts capturing *in vivo* binding that may include indirect recruitment via protein-protein interactions with other DNA-bound factors. Second, Wang et al. (2024) performed DAP-seq with recombinant *GmJAG1* protein capturing direct DNA-protein interactions *in vitro*.

ChIP-seq and DAP-seq peaks were mapped to genes based on genomic coordinates relative to transcription start sites (TSS). Peaks were classified by genomic location following the region definition used in Huang et al. (2021) as: promoter (-3000 bp to TSS), genic (TSS to transcription end site, TES) and downstream (+1 to +5000 bp from TES). This extended window captures binding in gene body regions beyond the canonical promoter. For each differentially expressed *JAG1* target, binding evidence was categorized as either “ChIP-seq only”, “DAP-seq only”, “both methods” or “no binding evidence”. Three experimentally validated *GmJAG1* DNA-binding motifs (nomenclature following Wang et al., 2024) were examined in candidate target gene promoters: M0 (GTTGGA), identified from ChIP-seq (Huang et al., 2021), and M1 (ACGCCACT) and M2 (ACTGGCAG), identified from DAP-seq (Wang et al., 2024).

### Weighted Gene Co-expression Network Analysis

TMM-normalized counts were transformed to log_2_-counts per million (logCPM) using the voom function (Law et al., 2014) for input to co-expression analysis. Weighted gene co-expression network analysis (WGCNA) was performed using the WGCNA R package (Langfelder and Horvath, 2008) to identify modules of co-expressed genes and relate them to leaf phenotypes. Genes were pre-filtered for significant expression variation using a moderated F-test (limma; Law et al., 2014) across all 10 experimental groups (2 leaf types x 5 timepoints). Genes with FDR < 0.01 were retained for network construction. A signed hybrid network was constructed using biweight midcorrelation (bicor), which considers both positive and negative correlations while focusing on positive connections within modules. The soft-thresholding power was selected as the lowest value achieving scale-free topology fit (R^2^ > 0.80).

Network construction used blockwise Modules with the parameters described above (full settings in analysis scripts). Module eigengenes (first principal component of each module) were calculated and correlated with line specific length-to-width (L:W) ratios (see above) using Pearson correlation. Fisher’s exact test was used to assess enrichment of *JAG1* candidate target genes within each module.

### Functional and Pathway Enrichment Analysis

Gene Ontology (GO) enrichment analysis was performed using clusterProfiler (Wu et al., 2021) with GO annotations for *Glycine max* obtained from Phytozome. Enrichment was tested separately for biological process (BP), molecular function (MF) and cellular component (CC) ontologies using Fisher’s exact test. For all enrichment analyses, the background gene set consisted of all expressed genes with GO annotations (see Results). Significantly enriched terms were identified as FDR < 0.05 with a minimum gene set size of 5 genes per term. GO enrichment was also performed on individual WGCNA modules to characterize module-specific biological functions.

Similarly, individual hormone pathway enrichment analysis was carried out using the KEGG Plant Hormone Signal Transduction pathway (gmx04075). KEGG Ortholog (KO) identifiers were mapped to soybean gene IDs using annotations from Phytozome. Each of the eight hormone pathways (auxin, cytokinin, gibberellin, ABA, ethylene, brassinosteroid, jasmonic acid and salicylic acid) were tested individually. Three levels of enrichment testing were performed using Fisher’s exact test. The first tested hormone enrichment among all DE genes at TP1 (narrow versus broad) to see whether leaf shape-associated genes are enriched for hormone pathway components. The second tested hormone enrichment among *JAG1* candidate targets (all tiers combined, see tiers definition above) to see whether genes regulated by *JAG1* are enriched for hormone signaling. The third tested direction-specific enrichment among upregulated versus downregulated DE genes separately, to see whether hormone pathway genes show a preferential regulation direction. Expected counts were calculated based on the proportion of each hormone pathway among all expressed genes. Each pathway was tested individually using Fisher’s exact test.

### Identification of High-Confidence *GmJAG1* Target Genes

High-confidence *GmJAG1* candidate targets were identified by requiring all four types of evidence described above: (1) differential expression, (2) binding evidence from ChIP-seq and/or DAP-seq datasets, (3) significant correlation between gene expression and leaf L:W ratio (Spearman |r| > 0.5, FDR < 0.05), and (4) membership in a WGCNA co-expression module that significantly correlated with leaf L:W ratio (P < 0.05). These two expression-based criteria are not independent where criterion 3 tests individual gene-phenotype association while criterion 4 requires that the gene additionally belongs to a co-regulated network associated with leaf shape.

### Analysis of *JAG1* Repression Machinery and Canonical Targets

#### JAG1 Repression Machinery Availability

To determine whether the complete repression machinery of *JAG1* (TOPLESS/TPR co-repressors and their recruited HDAC complex) is available when *JAG1* is active, we analyzed expression of TPR co-repressors and HDAC complex genes at the shoot apex (TP1). 12 soybean TOPLESS-related (*TPR*) genes were identified based on study by Yu et al. (2025) who comprehensively characterized the TOPLESS gene family in soybean. These include orthologs of Arabidopsis *TPL* (*GmTPR1* and *GmTPR2*) and *TPR1*-4. Similarly, 15 HDAC complex genes were curated from the *Glycine max* Wm82.a6.v1 genome based on PANTHER/KOG annotations containing “histone deacetylase” keywords, followed by manual validation to exclude five misannotated genes (two IST1-like/ESCRT-III proteins, two Sm D3 spliceosomal proteins, and one IST1P/UBP protein). The curated family includes nine catalytic subunits (orthologs of Arabidopsis RPD3/HDA6, HDA15, HDAC1 and HDAC11) and six regulatory subunits (SAP18 and SAP30L).

#### Cell Cycle Gene Family Analysis

Three cell cycle gene families were systematically characterized in soybean: *KRP* inhibitors, cyclins and CDKs. For each family, genes were identified using complementary approaches: Pfam domain search in Phytozome annotation (Wm82.a6.v1) and either BLASTP-based ortholog search against Arabidopsis or keyword search in functional annotations. Genes annotated as CDK inhibitors were excluded from the cyclin and CDK analyses.

Nine *KRP* genes were identified based on Guo et al. (2023) and verified using Pfam domain PF02234 (CDK inhibitor domain) and BLASTP against seven Arabidopsis *KRP* proteins (*KRP1*-*KRP7*). Both methods identified the same nine genes.

Cyclins were identified using Pfam domains PF00134 (Cyclin N-terminal) and PF02984 (Cyclin C-terminal) and classified by type: D-type (G1 phase), A-type (S-phase), B-type (M phase), U-type (plant-specific), T-type (transcriptional regulation) and H-type (CDK-activating kinase partner).

CDK genes were identified by keyword search and classified using Arabidopsis ortholog mapping based on three functionally characterized queries: *CDKA;1* (AT3G48750), *CDKB1;1* (AT3G54180) and *CDKB2;2* (AT1G20930).

For each gene across all three families, we determined basal expression levels (CPM) across all samples and at TP1, *JAG1* candidate target status based on differential expression, binding evidence from published ChIP-seq and DAP-seq datasets, and log_2_ fold change between narrow and broad genotypes. Cyclins and KRPs showing both differential expression and binding evidence were classified as high-confidence *JAG1* cell cycle targets.

#### D-type Cyclin Phylogenetic Analysis

D-type cyclin protein sequences were collected from *G. max*, *A. thaliana*, *M. truncatula* and *O. sativa* from Phytozome, identified by BLASTP and filtered for cyclin Pfam domains (PF00134, PF02984). After removing redundant rice paralogs, 70 sequences were aligned using MAFFT v7.520 (Katoh and Standley, 2013), trimmed using trimAl v1.4.1 (Capella-Gutiérrez et al., 2009) and used to infer a maximum likelihood tree with IQ-TREE2 v2.2.2.6 (Minh et al., 2020) under the Q.plant+I+G4 model with 1,000 ultrafast bootstrap and SH-aLRT replicates. The tree was rooted on *O. sativa CYCD3* and visualized using *ggtree* (Yu et al., 2017).

#### Integration with Published Single-Cell Transcriptomic Data

To independently validate the spatial expression of *JAG1* pathway components, we analyzed published single-nucleus RNA-seq (snRNA-seq) data from the soybean shoot apical meristem (SAM) and developing leaf (Fan et al., 2025). Pre-processed count matrices (SAM.scRNA.h5ad and Leaf.scRNA.h5ad) were obtained from the CNGB STOmics database (accession STDS0000375; https://db.cngb.org/stomics/). Gene identifiers were converted from Wm82 (Glyma) to ZH13 (SoyZH13) nomenclature using the SoyBase pangene lookup table (Glycine.pan5.MKRS; https://www.soybase.org).

Two complementary analyses were performed. First, a hypothesis-driven analysis tested spatial co-localization of pathway genes across six families (JAG1/JAG2, TPR, HDAC complex, KRP, cyclin, CDK) with *JAG1* expression in the SAM. For each gene, two metrics were computed across all 16 SAM cell types: the percentage of cells within each cluster expressing the gene and the mean normalized expression across all cells in each cluster. Spearman rank correlations were computed between *JAG1* percent expression and each gene family’s mean percent expression across clusters. *JAG1* and *JAG2* expressions were additionally compared between SAM and leaf tissues. SAM-versus-leaf comparisons for *KRP*s used Mann-Whitney U tests. Second, a data-driven analysis mapped all 1,567 *JAG1* candidate targets onto SAM clusters, computing per-cluster mean percent expressing and mean normalized expression. Targets were scored separately by confidence tier (Tier 1, Tier 2, Tier 3). Spearman and Pearson correlations were computed between *JAG1* percent expression and aggregate target expression across clusters to test for the predicted inverse relationship under active repression.

### Statistical Analysis and Software

Statistical analyses were performed in R v4.5.2 (R Core Team, 2025). Key packages included: edgeR v4.6.3 for normalization and differential expression (Robinson et al., 2010; Chen et al., 2016); vegan v2.7-2 for PERMANOVA analysis (Oksanen et al., 2025); WGCNA v1.73 for co-expression network analysis (Langfelder and Horvath, 2008); clusterProfiler v4.16.0 for functional enrichment (Wu et al., 2021); tximport v1.36.1 for transcript quantification import (Soneson et al., 2015); and sva v3.56.0 for batch correction. Single-cell integration analyses were performed in Python v3.10.19 using scanpy v1.11.5 (Wolf et al., 2018) and scipy v1.15.3 (Virtanen et al., 2020). Multiple testing correction used the Benjamini-Hochberg FDR method throughout. Visualizations were created using ggplot2 v4.0.1 pheatmap v1.0.13 and ComplexHeatmap v2.24.1.

## Acknowledgements

This work was supported by the Realizing Increased Photosynthetic Efficiency (RIPE) project, funded by the Bill & Melinda Gates Foundation under grant 57248 to the University of Illinois. We thank the Roy J. Carver Biotechnology Center at the University of Illinois for RNA library preparation and sequencing, and the RIPE facility for controlled environment growth chambers.

## Author Contributions

BGT conceived the study, performed the experiments, analyzed the data, and wrote the manuscript. CK collected the phenotypic data from growth chambers. EAA conceived the study, supervised the research, and revised the manuscript.

## Data Availability Statement

Raw RNA-seq data have been deposited in the NCBI Gene Expression Omnibus (GEO) under accession number GSE317596. Processed data and analysis scripts are available at https://github.com/bgtamang/Soybean-leaf-timecourse-global-transcriptomics. Interactive data explorer: https://bgtamang.shinyapps.io/GmJAG1-Dashboard/

## Supporting Information

**Figure S1.**
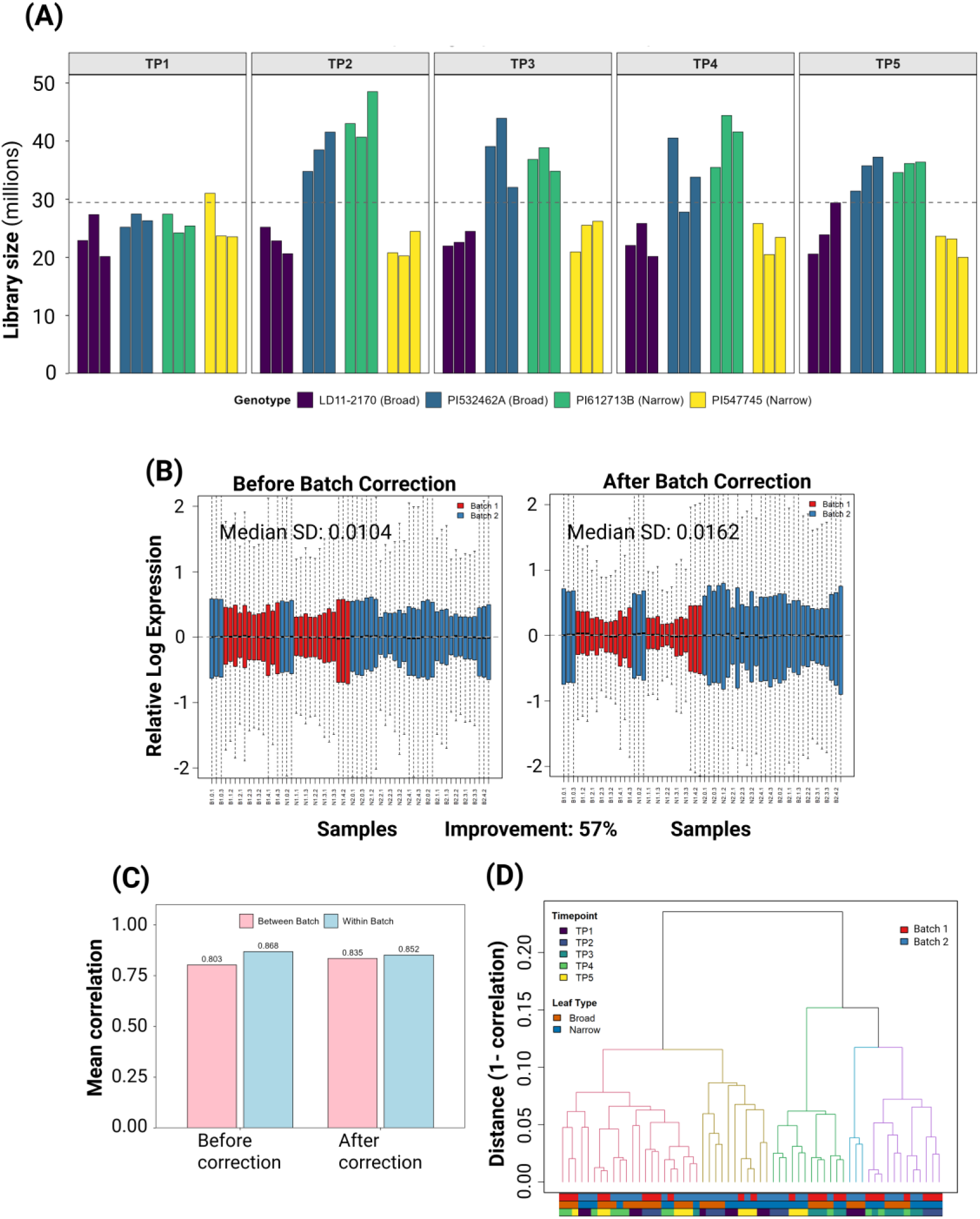
Extended RNA-seq quality control and batch correction assessment. **(A)** Library size (total sequencing reads) for all 60 RNA-seq samples, grouped by developmental timepoint (TP1-TP5). Within each timepoint, bars represent individual biological replicates for each of the four soybean genotypes (color-coded). The dashed horizontal line indicates the overall mean sequencing depth (29.5 million reads). **(B)** Relative Log Expression (RLE) plots before (left) and after (right) ComBat-seq batch correction. Each boxplot represents the distribution of expression deviations from the gene-wise median for one sample. Samples are colored by sequencing batch. The dashed horizontal line indicates the expected median of zero. **(C)** Comparison of mean sample correlations within and between sequencing batches before and after ComBat-seq correction. **(D)** Hierarchical clustering dendrogram of all RNA-seq samples after ComBat-seq batch correction. Samples were clustered using average linkage based on Pearson correlation distances (1- r). Colored bars below the dendrogram indicate developmental timepoint (TP1-TP5), leaf type (Broad, Narrow), and sequencing batch (1, 2). Samples cluster primarily by developmental stage rather than batch, indicating effective batch correction.

**Figure S2.**
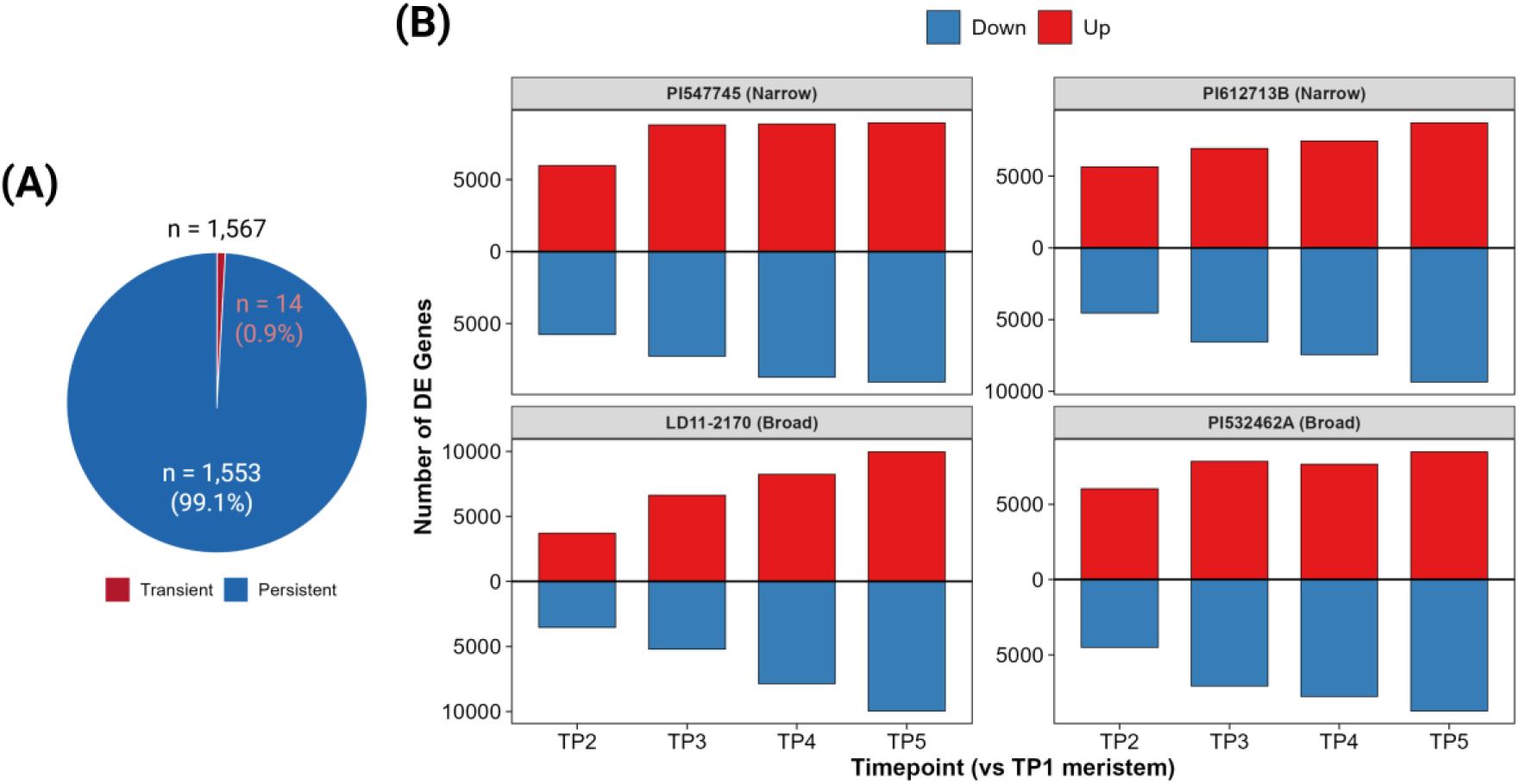
Temporal differential gene expression profiles by individual genotype. **(A)** Temporal expression persistence of *JAG1* targets. Pie chart showing that 99.1% of candidate targets (n = 1,553) maintained differential expression throughout development (blue), while only 0.9% (n = 14) showed expression differences restricted to early timepoints (red). **(B)** Diverging bar plots showing the number of differentially expressed genes at each developmental timepoint (TP2-TP5) relative to the shoot apex (TP1) for each of the four soybean genotypes used in this study. Two narrow-leaflet genotypes (PI 547745 and PI 612713B) and two broad-leaflet genotypes (LD11-2170 and PI 532462A) are shown. Red bars indicate upregulated genes and blue bars indicate downregulated genes (|log₂FC| > 1, FDR < 0.05).

**Figure S3.**
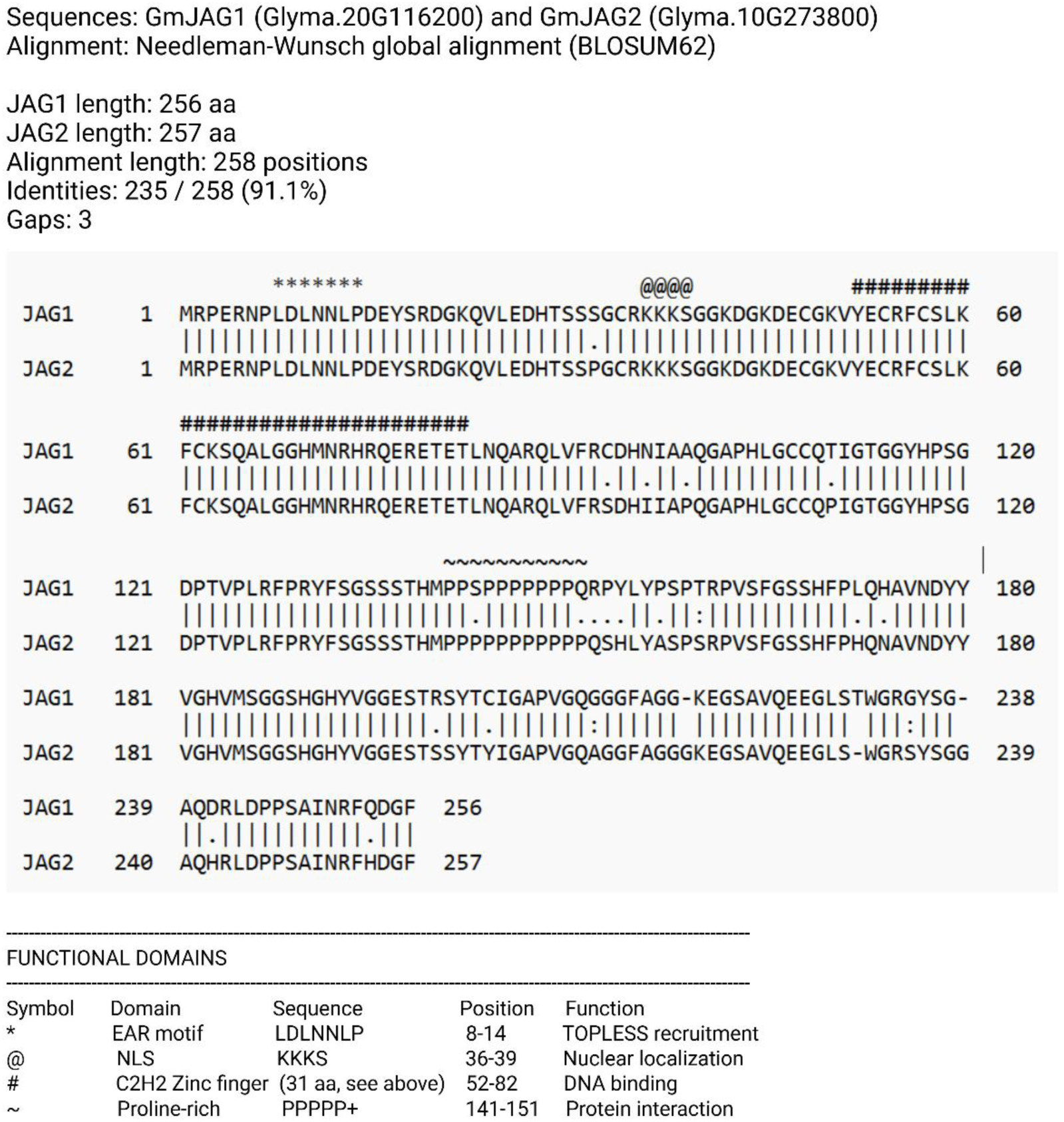
GmJAG1 and GmJAG2 protein sequence alignment. Pairwise alignment of GmJAG1 (Glyma.20G116200, 256 aa) and GmJAG2 (Glyma.10G273800, 257 aa) showing 91.1% amino acid identity. Protein sequences were translated from coding sequences deposited in GenBank: *JAG1* (JX119212.1) and *JAG2* (JX119214.1) from Jeong et al. (2012). Functional domains are annotated: EAR motif (*, position 8-14), nuclear localization signal (@, position 36-39), C2H2 zinc finger (#, position 52-82), and proline-rich region (∼, position 141-151). Both paralogs retain identical EAR motif (LDLNNLP) and C2H2 zinc finger domains including the QALGGH DNA-binding motif. Alignment performed using Needleman-Wunsch algorithm with BLOSUM62 matrix.

**Figure S4.**
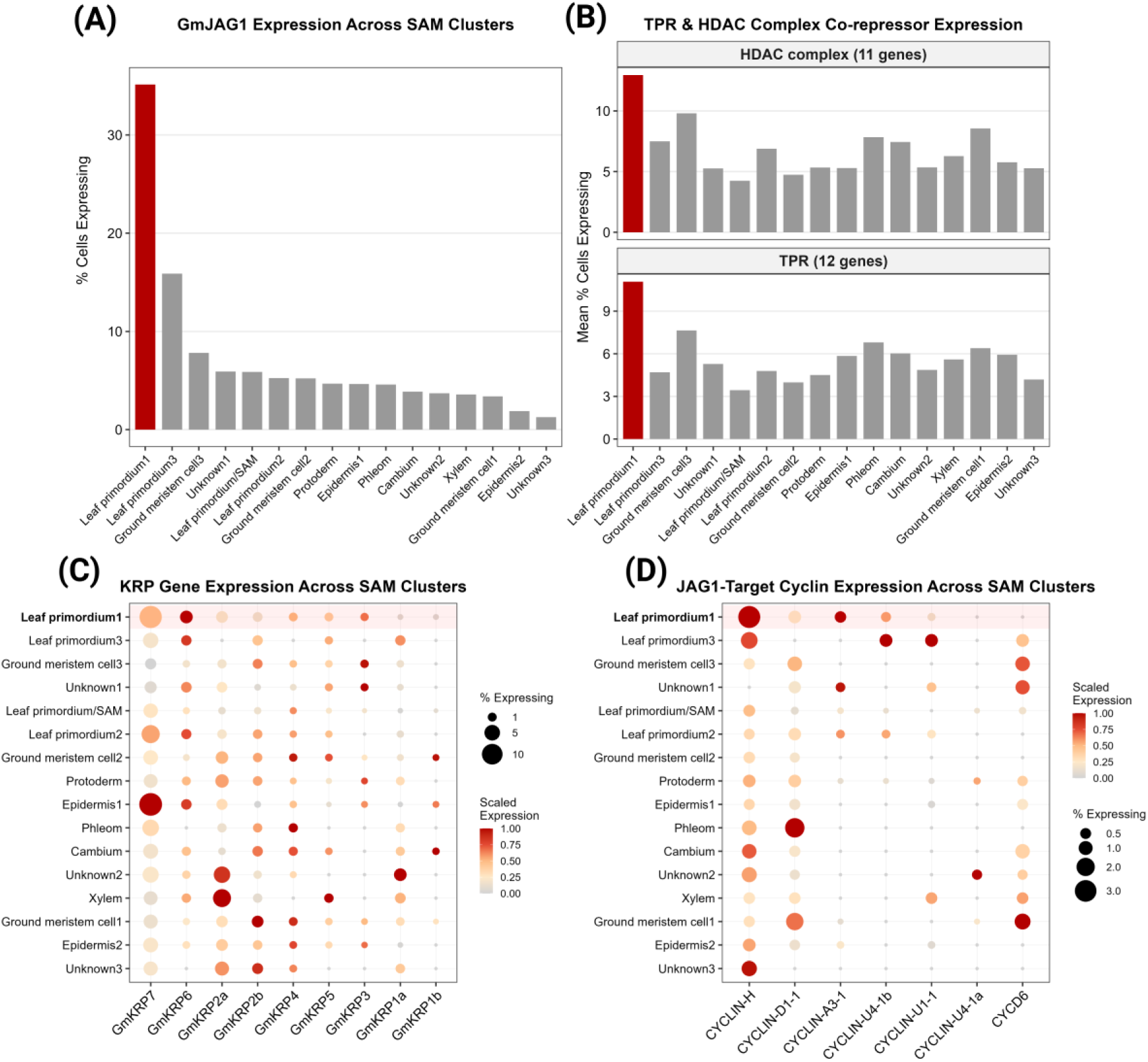
Single-cell expression of *JAG1* pathway components across SAM cell-type clusters. **(A)** GmJAG1 percent expressing across 16 SAM cell-type clusters from Fan, et al. (2025) snRNA-seq data (9,869 nuclei). LP1 = Leaf Primordium 1 (515 cells, red). **(B)** Mean percent expressing for *TPR* co-repressors (12 genes) and HDAC complex (11 genes) across all 16 SAM clusters. **(C)** Seurat-style dot plot of nine individual *KRP* genes across 16 clusters. Dot size represents percent of cells expressing; dot color represents per-gene scaled mean expression. (D) Seurat-style dot plot of seven *JAG1*-target cyclin genes across 16 clusters. Dot size and color encoding as in **(C)**.

**Figure S5.**
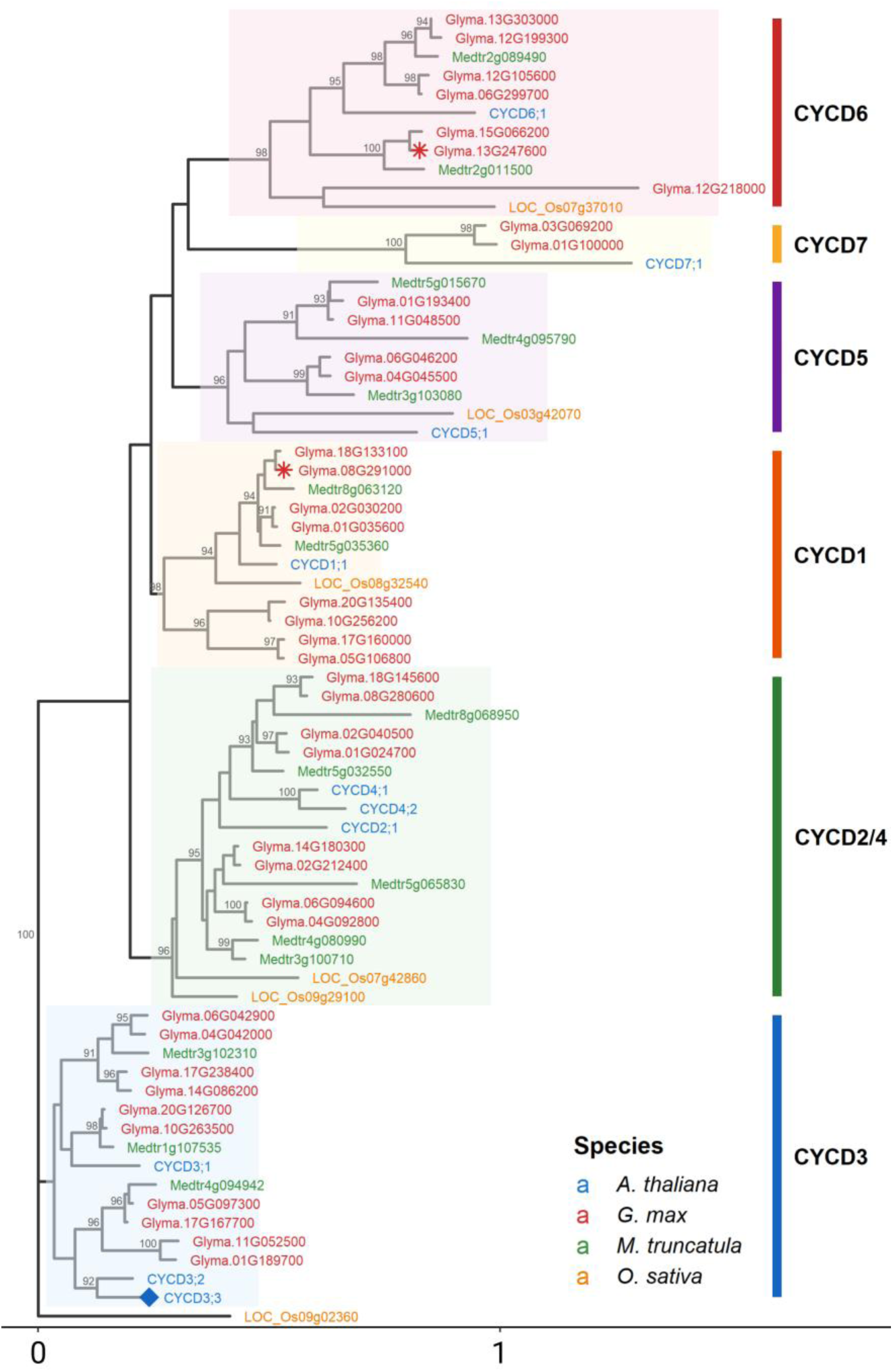
Phylogenetic analysis of D-type cyclins across four plant species. Maximum likelihood phylogeny of 70 D-type cyclin protein sequences from *Glycine max* (39 sequences), *Arabidopsis thaliana* (10), *Medicago truncatula* (15) and *Oryza sativa* (6 representatives, one per subclade). The tree is rooted on *O. sativa CYCD3*. Six conserved subclades (*CYCD1, CYCD2/4, CYCD3, CYCD5, CYCD6, CYCD7*) are indicated by colored background shading and labeled vertical bars. Red asterisks mark the two *GmJAG1*-regulated D-type cyclins in the *CYCD1* and *CYCD6* subclade. Blue diamond marks *AtCYCD3;3*, the Arabidopsis *JAG* target.

**Figure S6.**
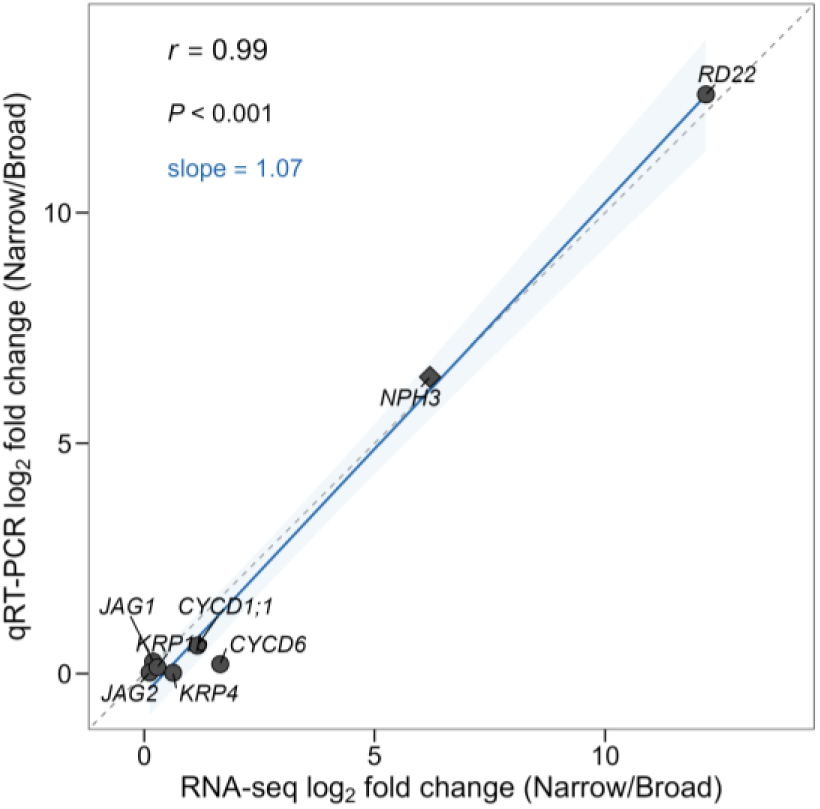
qRT-PCR validation of RNA-seq findings. Correlation between RNA-seq log₂ fold change and qRT-PCR log₂ fold change for eight genes validated across four soybean genotypes at TP1 (shoot apex). Each point represents the mean of two pairwise comparisons (PI 612713B vs PI 532462A and PI 547745 vs LD11-2170). Gene names are italicized. The dashed grey line indicates 1:1 identity; the blue line shows linear regression with 95% confidence interval. NPH3 (diamond) was undetectable by qPCR in PI 532462A; fold change was estimated using a Cq floor of 40. Pearson correlation coefficient, P-value, and regression slope are shown. Expression was normalized to the geometric mean of Actin 11 and RPL30 reference genes.

Table S1. Housekeeping gene expression stability validation.

Table S2. Complete list of 1,567 *JAG1* target genes with tier classification and expression data.

Table S3. Differential expression results for all genes across all comparisons.

Table S4. *KRP* gene analysis results with binding evidence.

Table S5. *KRP* expression analysis with CPM values.

Table S6. *KRP* trajectory analysis across timepoints.

Table S7. *TPR* and HDAC complex gene expression data.

Table S8. *TPR* trajectory analysis across timepoints.

Table S9. Gene Ontology enrichment results for all *JAG1* candidate targets.

Table S10. Gene Ontology enrichment results for Tier 1 candidate targets only.

Table S11. Hormone signaling genes by pathway with DE/*JAG1* status.

Table S12. Individual hormone pathway enrichment among *JAG1* candidate targets.

Table S13. Hormone enrichment by DE direction.

Table S14. Combined hormone enrichment summary across all analyses.

Table S15. Complete soybean cyclin gene family with Pfam domains and *JAG1* candidate target status.

Table S16. *JAG1*-targeted cyclin genes with binding evidence and expression patterns.

Table S17. Complete soybean CDK gene family with Pfam domains and *JAG1* target status.

Table S18. Cell cycle regulatory component comparison: CDKs vs Cyclins vs KRPs.

Table S19. Phenotype correlation analysis results.

Table S20. WGCNA module enrichment for *JAG1* candidate targets.

Table S21. Hub genes for each WGCNA module.

Table S22. WGCNA module correlations with phenotypic traits.

Table S23. Top 100 genes most strongly correlated with leaf L:W ratio.

Table S24. Binding evidence summary for all 1,567 *JAG1* candidate targets.

Table S25. Binding peak position analysis relative to transcription start sites.

Table S26. Integration of differential expression with *GmJAG1* binding data.

Table S27. Multi-evidence target classification.

Table S28. Top 100 priority targets ranked for functional validation.

Table S29. 79 high-confidence *GmJAG1* targets supported by all four evidence layers.

Table S30. Single-cell expression of *JAG1* pathway component families across 16 SAM clusters.

Table S31. *KRP* SAM-versus-leaf expression comparison.

Table S32. *JAG1* target co-localization across SAM clusters by confidence tier.

Table S33. qRT-PCR primer sequences and amplification efficiencies for validation of RNA-seq findings.

Table S34. qRT-PCR validation results for eight target genes across four soybean genotypes at TP1 (shoot apex).

